# Spatial Mapping of the Lung Cancer Ecosystem Reveals Distinct Patterns of Intratumoral and Internodular Heterogeneity

**DOI:** 10.64898/2026.08.31.748358

**Authors:** Bum-Seo Baek, Ki-Hyun Kim, Seung-jae Kim, Jacob D. Eccles, Minhyung Kim, Sungyong You, Christian Ascoli, Gye Young Park

## Abstract

The spatial organization of malignant and non-malignant cells within the tumor microenvironment (TME) critically influences tumor evolution and therapeutic response. However, the architecture of micro-niches remains incompletely understood. Leveraging Xenium-based spatial transcriptomics, we comprehensively mapped the spatial ecosystem of an orthotopic murine lung cancer model, identifying distinct spatial domains that form unique, organized cellular neighborhoods. These domains cluster into three major communities: (1) non-tumoral regions that recapitulate canonical normal lung structures; (2) a heterogeneous peri-tumoral region composed of spatial domains characterized by mesenchymal remodeling, active immune checkpoint signaling, and immunosuppressive myeloid populations; and (3) intra-tumoral regions that reveal marked tumor nodule heterogeneity, with unique tumor-specific domains exhibiting hallmark cancer pathways. Furthermore, our analytic approach was applicable to human lung cancer tissue. Notably, spatial domain analysis allowed us to resolve tumor nodules into multiple biologically distinct subtypes, defined by domain composition, hallmark cancer programs, and intercellular communication patterns within the TME.

**One Sentence Summary:** Spatial transcriptomics reveal diverse multicellular niches that organize intratumoral, peri-tumoral, and internodular heterogeneity in lung cancer.

## Introduction

Cancer cells and their neighboring non-cancerous structural or immune cells interact through both direct contact and secreted factors within unique tumor microenvironments (TMEs). These TMEs are further composed of micro-niches - distinct cellular neighborhoods where cancer cells, immune cells, and stromal cells assemble and communicate.(*1*)

The TME is a highly complex tumor ecosystem characterized by significant spatial heterogeneity, and the specific milieu is shaped by a wide variety of cellular and extracellular components.(*1*) These distinct microenvironments drive tumor diversity and play critical roles in metastasis and therapeutic resistance.(*2*) Micro-niche formation is influenced by factors such as anatomical structures, gradients of chemokines and cytokines, nutrient distribution, and cellular composition of the local environment. Furthermore, the various cell types within the TME undergo co-evolutionary adaptation in response to predominant niche factors, leading to distinct local neighborhoods that may be dissected based on their defining gene expression signatures.(*3*)

Advancements in spatial transcriptomics and analytical tools now enable us to investigate tumor ecosystems at high resolution, revealing the organization of micro-niches within the TME. Leveraging Xenium-based spatial transcriptomics, multicellular neighborhood analysis, and cell–cell communication profiling, we conducted a comprehensive characterization of these micro-niches using an orthotopic murine lung cancer model and applied this approach to human lung cancer. This approach identified distinct spatial domains and patterns of cellular interaction that define the organization of the tumor ecosystem. Furthermore, using a domain-based Cancer Index framework, we refined the delineation of the peri-tumoral region, characterized by prominent immune activity, and validated these classifications using orthogonal digital pathology. By integrating niche-level, domain-level, and tissue-level analyses, we demonstrate how local multicellular niches assemble into higher-order tumor structures and extend the characterization of tumor-associated microenvironments beyond previously described niche-level observations.(*4*)

## Results

### Xenium-based spatial transcriptomics unveils the multiscale architecture of the lung tumor ecosystem

To enable high-resolution spatial transcriptomic profiling of the lung cancer TME, we developed an integrated workflow that combines an orthotopic murine model with a high-plex Xenium platform (Fig. 1A). In this approach, we used our Scgb1a1-CreERT2;LSL-KP mice to generate primary lung cancer cells engineered to express a minimal neoantigen (minOVA), ensuring an appropriate immune response, as we previously reported.(*5*) After confirming stable minOVA expression, these cells were delivered intratracheally to three individual syngeneic recipients and allowed to progress for four weeks. Upon harvest, histopathological analysis identified representative regions spanning malignant and non-malignant states, providing essential context for subsequent spatial transcriptomic profiling.

**Fig. 1.**
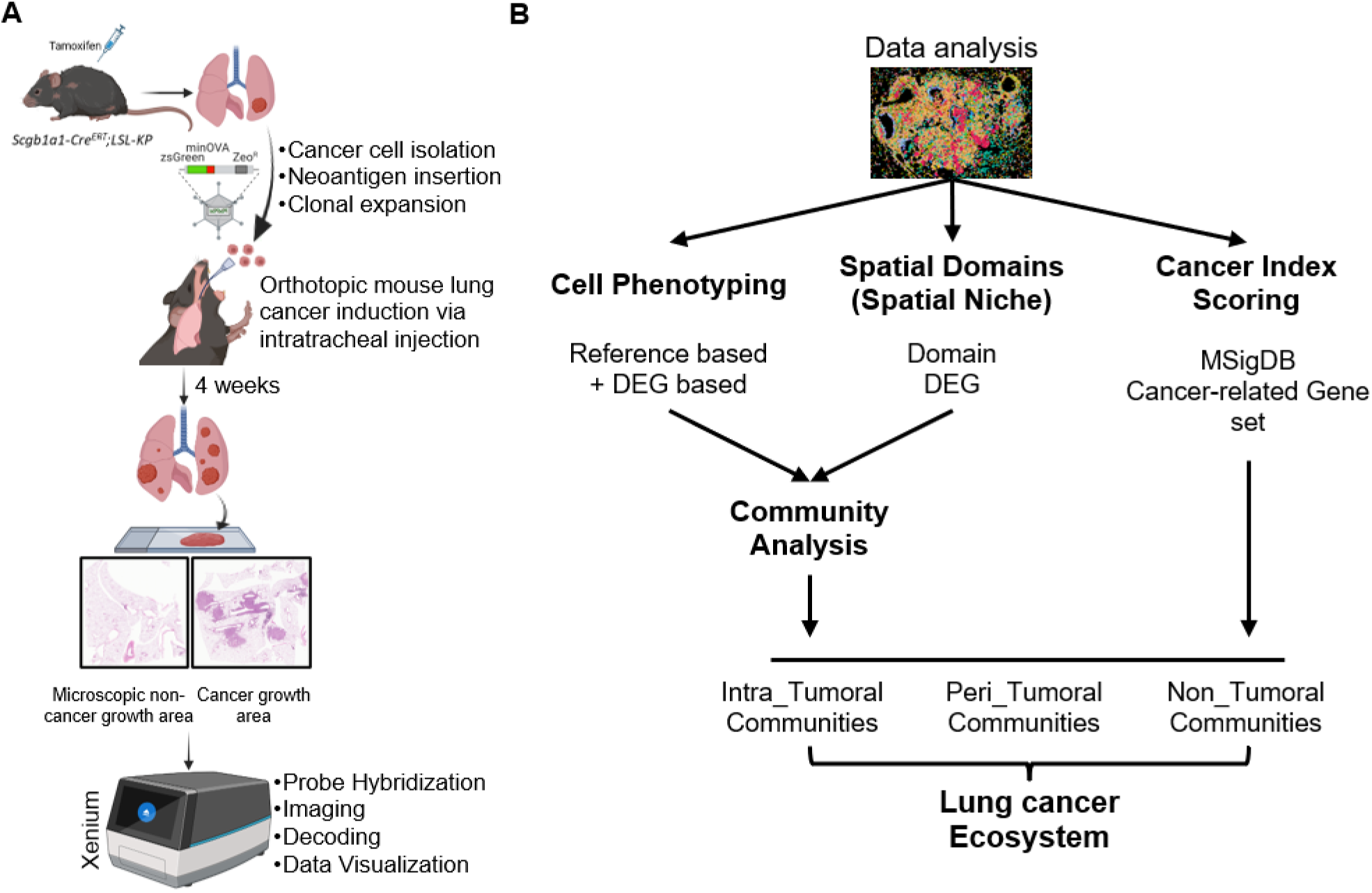
Spatial Transcriptomic Profiling of the Lung Tumor Ecosystem in an Orthotopic Murine Model. **(A)** Engineered lung cancer cells (Scgb1a1-CreERT2;LSL-KP derived, *zsGreen-minOVA*+) were orthotopically transplanted into syngeneic mice. Four weeks post-induction, tumors were analyzed using H&E histology and Xenium spatial transcriptomics. **(B)** Analytical pipeline: Following cell-type classification and BANKSY spatial clustering, domains were functionally annotated using Cancer Index and CellChat analysis. These domains were then stratified into intra-tumoral, peri-tumoral, and non-tumoral communities to define a comprehensive lung cancer spatial ecosystem.

The resulting multiplex transcript maps served as the foundation for spatial domain delineation using BANKSY clustering,(*6*) segmenting the tissue into distinct, spatially organized cell communities (Fig. 1B) that retained consistent transcriptional identities across tissue sections despite variation in abundance and spatial distribution among sections. By integrating reference-based cell classification using the Mouse Lung Cell Reference v1.1(Table S1)(*7*)-with a “Cancer Index” (a composite score derived from key oncogenic programs curated from MSigDB,(*8*) see Methods) and spatially constrained ligand-receptor analysis, we developed a multi-modal framework that systematically partitioned the tumor ecosystem into three hierarchical tiers comprised of non-tumoral, intra-tumoral, and a peri-tumoral interface regions.

### Characterization of spatial domain composition and regional features of tumors in lung cancer

We conducted a reference-based label transfer approach for cell-type annotation, based on the Mouse Lung Cell Reference v1.1. Assignments were further validated by assessing differentially expressed marker genes, enabling accurate classification across epithelial, endothelial, immune, and mesenchymal compartments. This dataset included a comprehensive range of lung-resident, infiltrating, and tumor-associated cellular populations within tumor-bearing lung tissue that formed well-defined clusters of major cell populations in UMAP space (Fig. S1), supporting the robustness of the annotation strategy and the presence of distinct transcriptional identities across our tissue sections. Epithelial, endothelial, and immune populations were the most abundant cell types, with major cell types (those representing more than 5% of the dataset) including CAP1/EPC (Capillary Endothelial Cell/Endothelial Progenitor Cell) cells (∼15%), alveolar epithelial cells (∼11%), B cells (∼9%), dendritic cells (∼8%), and interstitial macrophages (∼6%) (Fig. 2A, right / Fig. S2). Only 0.6% of all cells could not be confidently assigned to a reference cell type after annotation. These unassigned cells were retained for cell-type composition summaries but excluded from subsequent spatial analyses when appropriate.

**Fig. 2.**
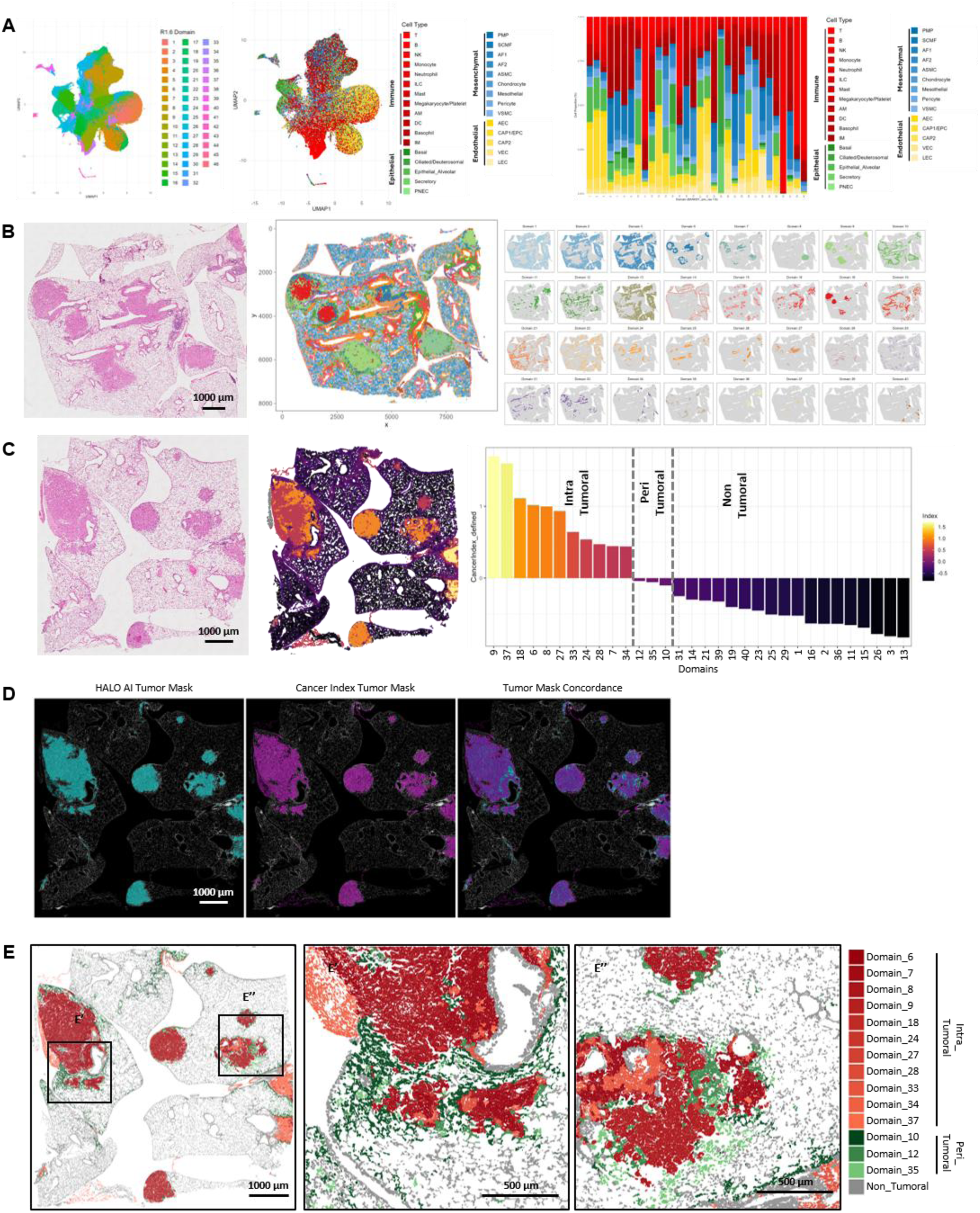
Spatially Resolved Multicellular Domains Define the Tumor Microenvironment. **(A)** UMAP embedding of the tumor microenvironment. Visualization of 535,796 cells colored by 46 spatial domains identified by BANKSY clustering (resolution 1.6, left) and reference-based cell types (right). **(B)** Multicellular domain composition. Stacked bar plots showing the proportion of cell types within each domain, revealing that spatial domains represent heterogeneous cellular admixtures rather than single cell types. **(C)** Spatial validation. Comparison of adjacent H&E histology with Xenium spatial domain maps. Individual domain highlight panels (right) confirm that BANKSY domains localize to specific histological structures. Functional stratification via Cancer Index. Domains were ranked by a “Cancer Index” derived from aggregated MSigDB hallmark gene set scores. This stratification categorizes domains into Intra-tumoral (high index), Peri-tumoral (intermediate), and Non-tumoral (low index) groups. **(D)** Representative spatial comparison of HALO-derived ground-truth tumor regions, BANKSY/Cancer Index–derived tumor classification, and their merged overlay in the 4w2 tissue section. **(E)** Ecosystem topology. Tissue-scale visualization showing the spatial organization of domain groups: Intra-tumoral (shades of red), Peri-tumoral (shades of green), and Non-tumoral (dark gray). Zoomed-in insets (E’, E’’) highlight the distinct boundaries between these microenvironments.

To identify recurrent multicellular microenvironments across tissue sections, we applied the BANKSY algorithm in a multi-sample integration setting, leveraging both transcriptional and spatial information to define shared spatial domains across independent tissue sections. For this, we selected a clustering resolution (Fig. S3) that captured biologically meaningful tissue organization consistent with canonical lung histology and confirmed that the resulting spatial organization was robust to parameter variation and reproducible across alternative spatial clustering approaches (Fig. S4). This analysis identified 46 distinct spatial domains (Fig. 2A, left) with conserved transcriptional identities that varied in abundance and spatial distribution between samples.

Spatially informed clustering using UMAP revealed a structured organization of domains that reflects both transcriptional similarity and spatial context across samples (Fig. 2A, left). Instead of forming isolated, discrete clusters, many domains occupied adjacent or partially overlapping regions on UMAP space, indicating graded relationships between spatial domains with related transcriptional programs. This continuous organization suggests that spatial domains capture transitions between microenvironmental states rather than strictly separated compartments, consistent with the spatial continuity of tumor-bearing lung tissue. Importantly, spatial domains were not composed of single cell types; instead, they represented multicellular admixtures, as demonstrated by the distribution of cell types across domain assignments in UMAP space (Fig. 2A, middle and right). This finding suggests that spatially informed clustering identifies coordinated neighborhoods of transcriptionally complementary cells rather than discrete lineage-based compartments, motivating further examination of domain composition and dominance.

Analysis of spatial domains across individual tissue sections revealed that reduced-dimensional organization captures common structural features of the tumor microenvironment, although section-specific differences exist in the prevalence and arrangement of spatial niches (Fig. S5).

Mapping BANKSY domains back to tissue coordinates revealed that many domains form spatially discrete, recurrent structures rather than being diffusely intermingled (Fig. 2B). Domain-specific panels further highlight that individual domains localize to specific tissue microenvironments, and direct comparison with matched H&E images from the same tissue sections confirmed concordance between inferred spatial domains and underlying histologic architecture. These results reinforce that BANKSY captures meaningful spatial organization within the tumor section.

To objectively annotate the tumor and its surrounding regions (Fig. 2C, left), we computed a domain-level Cancer Index by aggregating Hallmark module scores (EMT, Hypoxia, KRAS_DN, MYC V1/V2, apoptosis, TNFA–NFκB, IL6–JAK–STAT3, Glycolysis) from the Molecular Signatures Database (MSigDB) for each domain (Fig. S6), followed by z-scoring, directional alignment, and equal-weight averaging (Fig. 2C, middle and right).(*8*) Ranking domains by the Cancer Index revealed a clear gradient from high to low values, allowing stratification into intra-tumoral, peri-tumoral, and non-tumoral groups based on index distribution and spatial context. Using this approach, domains were annotated as intra-tumoral (Domains 6, 7, 8, 9, 18, 24, 27, 28, 33, 34, and 37), peri-tumoral (Domains 10, 12, and 35), with all remaining domains classified as non-tumoral (Fig. 2C, right, Table S2).

To validate this regional classification derived from transcriptomics, we compared tumor regions defined by our Cancer Index–based domain classifications with unbiased independent digital pathology annotations (AI-assisted HALO). Overall, Cancer Index–based domain classifications showed strong concordance with HALO-based tumor annotations, achieving balanced accuracies of 0.867 with high Matthews correlation coefficients (Fig. 2D, Fig. S7). Transcriptomics and pathology-derived tumor concordance was further supported by strong spatial overlap between the methods, as reflected by Dice coefficients of 0.767 and 0.779. Furthermore, tumor burden estimates were highly comparable, with Xenium-to-HALO tumor area ratios of 0.97 and 0.93, demonstrating that transcriptomics-derived classifications accurately recapitulated the overall tumor extent identified by pathology, supporting robust agreement between morphology-based and transcriptomics-based approaches to tumor annotation.

Having established the validity of Cancer Index–based domain classification, we next examined the spatial organization of these domain groups across entire tissue sections. Tissue-wide visualization of these grouped domain assignments showed that intra-tumoral regions occupy large, dense tumor patches, while peri-tumoral regions localize along tumor boundaries and adjacent transition zones. Non-tumoral regions broadly correspond to histologically normal areas (Fig. 2E). Zoomed-in regions of interest further revealed sharp yet locally mixed interfaces between intra-and peri-tumoral states, supporting that Cancer Index-guided domain grouping reconstructs a coherent tumor–adjacent–normal spatial axis across the section (Fig. 2E’, 2E’’).

### Identification and validation of non-tumoral structural domains through analysis of spatial domains

To systematically characterize lung architecture in non-tumoral areas, a comprehensive analysis was conducted on spatial domains annotated as non-tumoral (Fig. 2D). The analysis focused on tissue structure, cell-to-cell communication networks, and pathway activities within these spatial domains. Enriched pathways analysis identified coherent clusters of non-tumoral spatial domains with shared signaling profiles (Fig. 3A). These clusters were further resolved into distinct structures corresponding to the mesothelium (Domain 14), blood vessels (Domains 26, and 31), airways (Domains 15, 23, 25, and 29), parenchymal lung (Domains 1, 2, 3, 21, and 13), bronchus-associated lymphoid tissue (BALT; Domains 36, 39, and 41), and peri-BALT(Domains 11, 16, and 19) (Fig. 3A, 3B). This approach effectively recapitulated normal lung structural communities, each comprising multiple distinct domains.

**Fig. 3.**
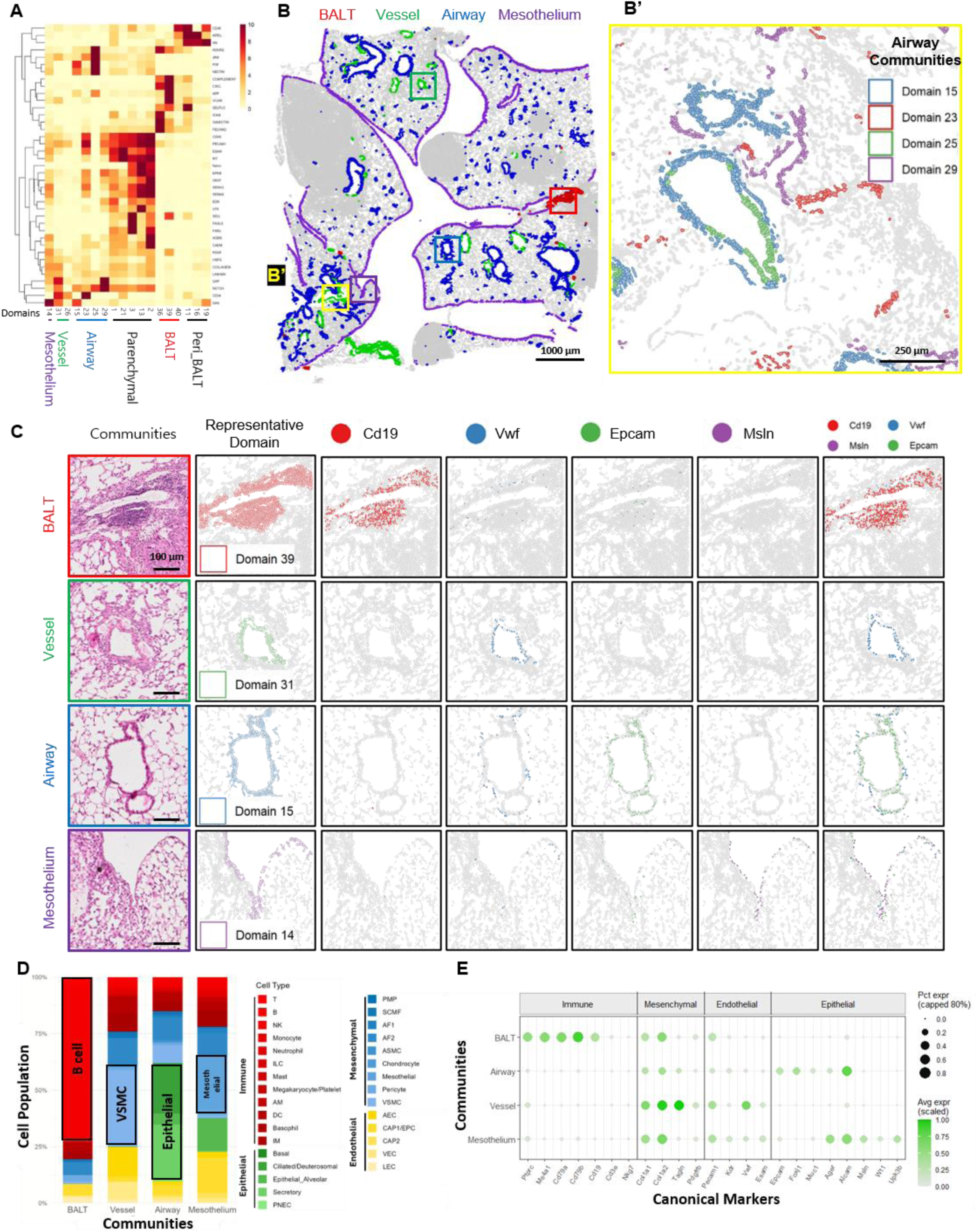
Identification and validation of non-tumoral structural communities through analysis of spatial domains and cell-to-cell communication networks within these domains. **(A)** Identification of structural communities: A CellChat pathway information flow heatmap reveals distinct signaling patterns among non-tumoral domains, clustering them into structural communities: mesothelium, vessel, airway, parenchymal lung, and BALT (Bronchus-Associated Lymphoid Tissue). **(B)** Spatial visualization of these communities: (Left) Xenium map highlights BALT (red), vessel (green), airway (blue), and mesothelium (purple) communities against a gray background. (Right) Magnified view (B’) detailing the organization of airway-specific domains (Domains 15, 23, 25, 29). **(C)** Structural validation: Adjacent H&E histology (left) is compared with the dominant spatial domain (middle) for each community - BALT (D39), vessel (D31), airway (D15), and mesothelium (D14) - while spatial gene expression plots (right) visualize the domain-restricted expression of canonical markers (*Cd19* for BALT, *Vwf* for vessel, *Epcam* for airway, *Msln* for mesothelium). **(D)** Cellular composition: Stacked bar plots quantify cell type proportions within each domains, confirming enrichment of B cells in BALT, VSMCs in vessels, epithelial cells in airways, and mesothelial cells in the mesothelium. **(E)** Canonical marker validation: A dot plot summarizes the percentage and average expression of lineage-specific genes across the representative structural domains, further validating their biological identities.

The clustered non-tumoral spatial domains were mapped onto lung tissue to provide spatial context and were consolidated into major structural communities (Fig. 3B). This mapping demonstrated that spatial domains are consistently localized to canonical lung structures across non-tumoral regions. Notably, the composite of airway spatial domains recapitulated the architecture of both large and small airways, while preserving fine-scale, contiguous, and domain-specific segments along airway structures (Fig. 3B’). These findings support the presence of structured airway niches composed of multiple, spatially interweaved micro-niches, rather than a single uniform airway structure. Collectively, these data show that our spatial domains robustly capture reproducible, functionally, and anatomically distinct lung structural communities, consistent with established lung histology and biology.

Each structural community features a representative domain, alongside additional supporting domains; for instance, airway domain 15 corresponds to a Foxj1^+^ ciliated airway epithelium niche, blood vessel domain 31 marks a perivascular-smooth muscle-pericyte niche, and BALT domain 39 represents a CD19-enriched B-cell niche, domain 14 identifies a pleural mesothelium-serosal lining niche (Fig. 3, Table S2). Matched H&E images were used to provide morphological context for representative domains, validating their spatial assignments and canonical marker expression patterns (Fig. 3C). Notably, the spatial expression patterns of canonical markers closely corresponded to established identities: *Cd19* is enriched in the BALT domain (Domain 39), *Vwf* in the vascular domain (Domain 31), *Epcam* in the airway epithelial domain (Domain 15), and *Msln* in the mesothelial domain (Domain 14), with each marker spatially concentrated within its respective domain.

Consistently, the cell-type composition within each structural community further supported their assigned identities (Fig. 3D). The BALT domain was enriched for B cells, the vessel domain for vascular smooth muscle cells (VSMCs), the airway domain for epithelial cells, and the mesothelium domain for mesothelial cells. These results suggest that cellular composition generally aligns with the established anatomical communities.

Finally, we assessed the prevalence and intensity of canonical community-defining genes across representative domains (Fig. 3E). The BALT displayed enrichment for B-cell markers (e.g., *Cd19/Ms4a1/Cd79a/b*), blood vessel domains for vascular/smooth muscle programs (e.g., *Vwf/Acta2/Tagln*), airway domains for epithelial/airway markers (e.g., *Epcam, keratins, Foxj1, Scgb1a1*), and the mesothelium for mesothelial markers (e.g., *Msln, Wt1, Upk3b*). These findings are consistent with the known active pathways in lung structural communities.(*9*)

Collectively, our spatial domain analysis in non-tumoral regions accurately recapitulates established lung structure and function, underscoring the potential to apply this approach to tumor-involved lung areas.

### Dissecting tumoral heterogeneity through spatial domains and signaling signature analysis

Several spatial domains are found exclusively within intra-tumoral regions, highlighting the unique characteristics of tumor-specific micro-niches (Fig. 2D). The defining features and characteristics of these domains are detailed in Table S2. By treating these tumor-specific spatial domains (micro-niches) as distinct functional units, we deconstructed tumor nodules into composites of these domains and identified characteristic types across our samples (Fig. 4A). Type 1 nodules were noted to be round and located in the peripheral lung field, attached to the pleura, and characterized by a high proportion of spatial domain 8, which is identified as a myeloid cell-like metaplastic stroma (Table S2). Type 2 nodules were situated in the central lung parenchyma and were found to be dominated by domain 7, which represents a CXCL9-high inflammatory CAF (Cancer Associated Fibroblast) niche. Type 3 nodules were polygonal, adjacent to the airways, and enriched for domain 9, featuring a chondrocyte stroma niche (Table S2). Type 4 nodules were also airway-adjacent, with a high representation of domain 18, defined by inflammatory response and ECM-anchoring stroma. Type 5 nodules were located near blood vessels and enriched for domain 37, which corresponds to proliferating interface progenitors. Contrastingly, mixed nodules lacked a dominant domain and instead exhibited a heterogeneous distribution of spatial domains (Fig. 4A), highlighting the spectrum of multicellular organization present across tumor nodules. Consistent with these observations, comparison of intra-tumoral domain composition across nodules revealed that each nodule type was defined by a characteristic combination of spatial domains rather than by a single domain alone (Fig. 4B).

**Fig. 4.**
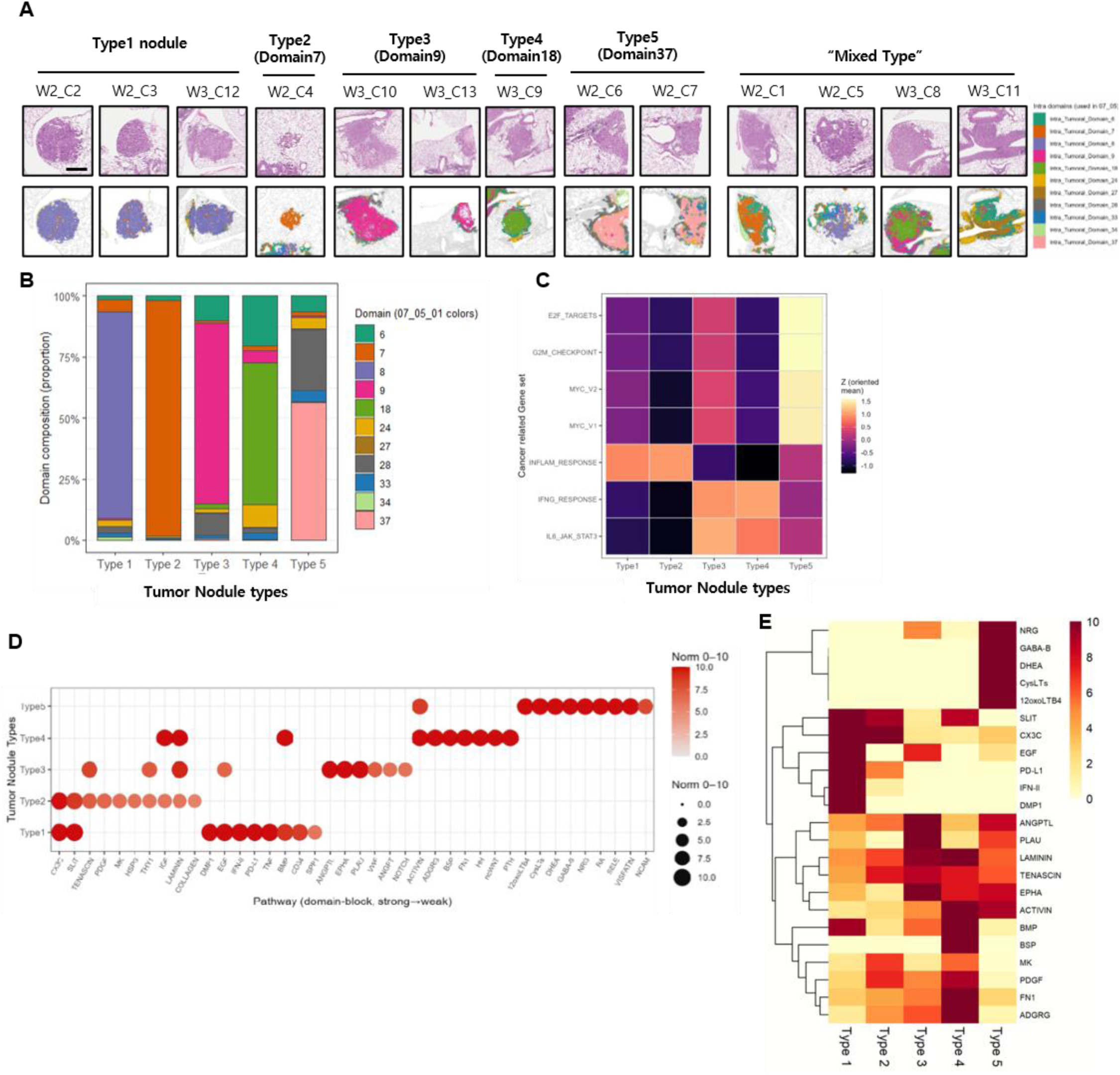
Dissecting tumoral heterogeneity through spatial domains and signaling signature analysis. **(A)** Tumor nodule classification: Tumor nodules were classified into five types (Types 1–5) based on their dominant intra-tumoral spatial domains. Representative H&E images (top) and spatial domain maps (bottom) highlight the distinct architecture of each nodule type. **(B)** Nodular heterogeneity with distinct spatial domain composition: A stacked bar plot quantifies the proportional composition of intra-tumoral spatial domains within each nodule type. **(C)** Cancer hallmark analysis: Heatmap of Z-scored hallmark gene set signatures across nodule types demonstrates that Type 5 is enriched for proliferative programs (E2F, G2M, MYC targets), whereas Types 1 and 2 show robust expression of inflammatory response signatures. Types 3 & 4 are characterized by strong IL6-JAK-STAT3 and IFN-γ signaling profiles. **(D)** Divergent signaling networks among nodule types: A dot plot displays the top 10 CellChat signaling pathways identified in each tumor nodule type, revealing distinct communication landscapes. **(E)** Type-specific molecular features: A heatmap shows the normalized expression of key functional genes (e.g., *Cd274* [PD-L1], *Egf, Fn1, Plau*) that differentiate the biological identities of the five nodule types.

Next, we investigated whether each domain-defined nodule type displayed distinct cancer-related transcriptional states. Hallmark gene sets at single-cell resolution have been used to analyze biological processes that reflect cancer-like phenotypes.(*8*) Our data show that Hallmark gene sets exhibited distinct expression patterns among the different domain-based tumor nodule types (Fig. 4C). Proliferation-associated signatures such as E2F targets, G2M checkpoint, and MYC targets were preferentially elevated in type 5, consistent with previously described cell cycle–high tumor states.(*10, 11*) In contrast, inflammatory and cytokine-related programs, including inflammatory response (Types 1 & 2), IFNγ response, and IL6/JAK/STAT3 signaling (Types 3 & 4), were distinctly enriched in other types. This suggests that immune engagement and inflammatory signaling are organized into specific, type-dependent nodule states rather than being uniformly distributed, although further validation is needed.

Lastly, we analyzed intercellular communication across the five tumor types and observed marked differences in dominant signaling pathways among nodule types (Fig. 4D). Type 1 and Type 2 nodules showed relatively stronger enrichment of inflammatory and cytokine-associated pathways (e.g., CX3C, IFN-II, PD-L1), whereas Type 3 and Type 4 nodules were characterized by increased activity of extracellular matrix– and adhesion-related signaling, including FN1, LAMININ, TENASCIN, and integrin-associated pathways (Fig. 4D, 4E). Type 5 nodules exhibited a distinct pattern with comparatively higher activation of neuroactive and lipid-associated signaling modules (e.g., NRG, GABA-B, CysLTs). Notably, hierarchical clustering of pathway activity scores distinguished tumor types based on their signaling profiles, suggesting structured differences in potential intercellular communication patterns across nodule architectures. (Fig. 4E). Although the sample size is limited, these findings demonstrate that this analytic approach can provide valuable insights into the functional characteristics of individual tumor nodules.

### Distinct tumor-specific spatial domains: mesothelial-enriched and lymphatic endothelial cell micro-niches

Spatial domain analysis revealed a unique mesothelial micro-niche abutting the tumor nodule (tumor-associated mesothelium; Domain 34), which was both compositionally and transcriptionally distinct from non-tumoral mesothelium (Domain 14). These two domains form a continuum along the pleural lining and are enriched in mesenchymal-lineage populations (characterized by the expression of Wt1, Msln, and *Upk3b*) (Fig. 5A-C, Table S3). However, tumor-associated mesothelium displayed a greater contribution of stromal and fibroblast-like subsets such as SCMF (secondary crest myofibroblasts) cells, whereas the non-tumoral mesothelium contained relatively more endothelial and minor epithelial components (Fig. 5A). Differential expression analysis revealed strong upregulation of immune-and stress-associated genes (e.g., *C1qa, C1qb, C3ar1, Angptl4, Dusp5*) in tumor-associated mesothelium (Fig. 5B), consistent with previous reports of cancer-associated mesothelial cells.(*12*) This suggests that mesothelial cells adjacent to tumor tissue adopt a remodeled inflammation-associated state. Spatial mapping of these mesothelial domains onto tissue sections revealed that tumor-associated mesothelium is confined to a narrow, contiguous interface bordering tumor regions. This area is characterized by preferential expression of *Angptl4*-and *Dusp5*-positive cells within the tumor-adjacent mesothelial band, whereas non-tumoral (normal) mesothelium remains distributed along canonical pleural surfaces.(Fig. 5C-C’) Spatial mapping of mesothelial cell-to-cell communication within these domains revealed distinct matrix-adhesion signaling networks such as *Fn1-Sdc1* and *Col1a1-Sdc1* in tumor-adjacent mesothelium compared with non-tumoral mesothelial tissue (Fig. 5D), suggesting active crosstalk between mesothelial cells and neighboring structural cells mediated by extracellular matrix signaling. Spatial imaging further visualized these features, highlighting spatially patterned expressions of *Col1a1, Col1a2, and Sdc1* in tumor-associated mesothelium (Fig. 5E).(*13, 14*)

**Fig. 5.**
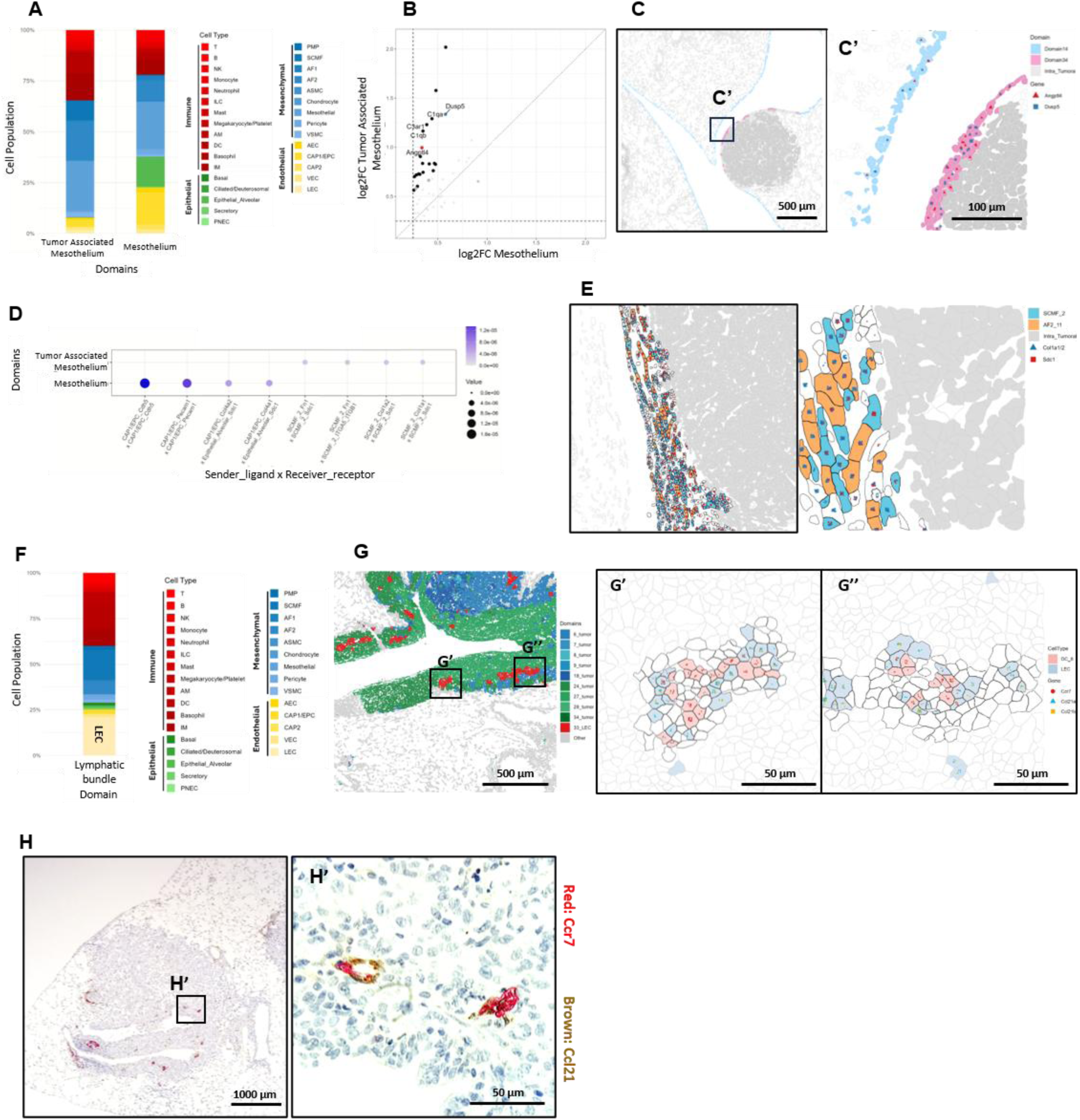
Distinct tumor-specific spatial domains: mesothelial-enriched and lymphatic endothelial cell micro-niches. **(A)** Stacked bar plots compare the cellular composition of tumor-associated (D34) and non-tumoral (D14) mesothelium, demonstrating a marked expansion of stromal SCMF populations in tumor-associated mesothelium. **(B)** Differential expression analysis indicates that tumor-associated mesothelium is characterized by upregulation of immune- and stress-response genes such as *C1qa, C1qb, Angptl4, Dusp5*. **(C)** Visualization of the pleural boundary shows a spatial continuum and transition from non-tumoral to tumor-associated mesothelium. **(D)** Matrix-adhesion signals are enriched in tumor-associated mesothelium. **(E)** High-resolution spatial mapping of tumor-associated mesothelium (D34) shows spatially patterned expression of matrix remodeling markers (*Col1a1, Col1a2, Sdc1*) at the tumor interface. **(F)** Domain 33 represents a lymphatic endothelial micro-niche, with a cellular composition highly enriched in lymphatic endothelial cells (LECs). **(G)** Spatial visualization of the LEC-enriched domain (D33) demonstrates co-expression of the *Ccl21-Ccr7* signaling axis, consistent with the canonical lymphatic function in immune cell trafficking. **(H)** Immunohistochemical validation of CCR7–CCL21 signaling in the tumor-bearing lung section. Low-magnification image shows Ccr7-and Ccl21-positive structures within the tumor region, with the boxed area indicating the region shown in H′. **(H′)** Magnified view of the boxed region in H, showing Ccr7-positive and Ccl21-positive signals localized to vascular-like structures within the tumor microenvironment, supporting the presence of an LEC-associated lymphatic chemotactic niche.

In addition, we identified another distinct tumor-specific spatial domain (Domain 33) within the intratumoral region that was highly enriched in lymphatic endothelial cells (LECs) (Fig. 5F-G). This LEC-rich micro-niche was characterized by strong expression of LEC-specific markers such as *Ccl21a, Ccl21b, and Prox1* (Table S3), consistent with previous findings.(*15*) Previous studies have shown that tumor-associated lymphatic vessels play pleiotropic roles in immune cell trafficking, metastatic dissemination, and shaping the local immune microenvironment.(*16*) In our analysis, the spatially defined LEC-enriched domain formed a hollow structure and displayed a cell–cell communication pattern dominated by CCR7–CCL21 signaling, consistent with a canonical lymphatic chemotactic program (Fig. 5G, 5G’,5G’’).(*17, 18*)

To further validate the presence of intratumoral LEC structures (Domain 33) at the protein level, we performed immunohistochemical (IHC) staining for CCR7 and CCL21 on tissue sections of the remaining samples. In these tissue sections, clusters of CCR7-and CCL21-positive cells were predominantly observed within tumor-involved areas, and higher-magnification imaging confirmed their architectural organization into vessel-like tubular structures, resembling the transcriptomics-defined LEC domains (Fig. 5H, H′). Together, these results suggest that our domain-based approach enables high-resolution mapping of intratumoral communities and facilitates the identification of tumor-specific communities.

### Spatial domains at the tumor boundary reveal distinct stromal, immune-active, and suppressive myeloid niches

A distinct set of spatial domains (Domains 10, 12, and 35) was found to be concentrated at the interface between the intra-tumoral core and the adjacent normal tissue, forming a heterogeneous rim-like peri-tumoral region (Fig. 2D, 2E). Rather than constituting a uniform boundary, this peri-tumoral compartment comprised several distinct micro-niches, each characterized by unique cellular compositions, transcriptional signatures, and patterns of intercellular communication (Fig. 6B–D).

**Fig. 6.**
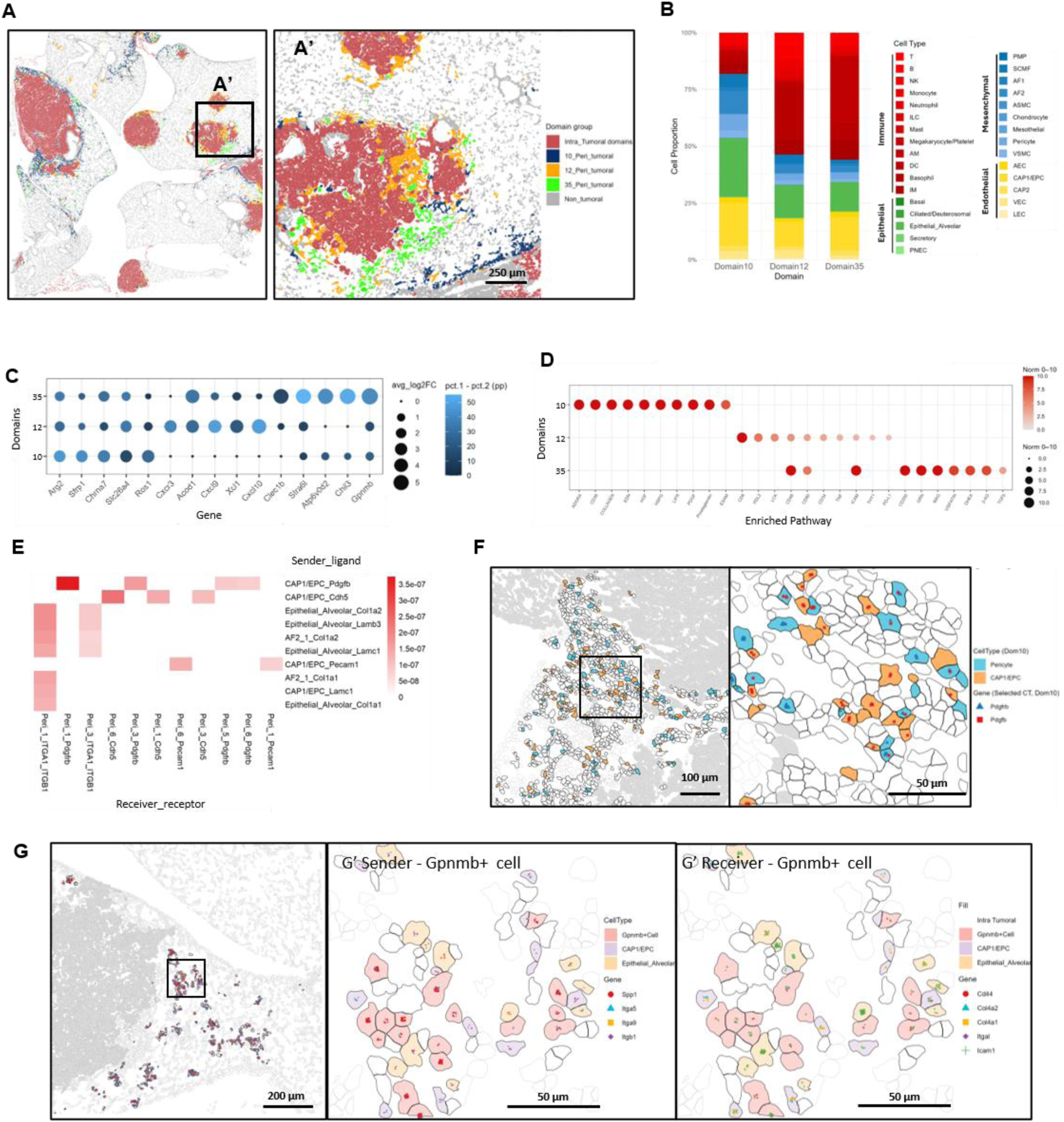
Spatial domains at the tumor boundary reveal distinct stromal, immune-active, and suppressive myeloid niches. **(A)** Heterogeneous rim organization at the tumor boundary: Spatial mapping visualizes distinct micro-niches (Domains 10, 12, and 35) specially localized at the interface between the intra-tumoral core and adjacent non-tumoral tissue. **(B)** Peri-tumoral domain composition: Domain 10 is enriched with mesenchymal cells, while Domains 12 and 35 are predominantly composed of immune cells. **(C)** Domain-specific transcriptional signatures: Domain 10 displays strong stromal remodeling markers (e.g., *Sfrp1)*, whereas Domain 12 is characterized by expression of T-cell recruiting chemokines (*Cxcl9, Cxcl10, Xcl1*). Domain 35 exhibits M2-like/LAM signatures (*Gpnmb, Chil3*). **(D)** CellChat pathway analysis: Matrix/PDGF signaling is prominent in Domain 10, while Domain 35 shows a robust CD200/TGF-β signaling activity. **(E)** Ligand-receptor interaction map of domain 10. **(F)** Spatial distribution of mesenchymal cells (pericytes) and epithelial cells across the tissue section highlights the proximity of *Pdgfb-*and *Pdgfrb-*expressing cells. **(G)** High-resolution spatial visualization of Domain 35 shows the close proximity of *Gpnmb*-expressing macrophages to cells expressing *Spp1,* Integrin *Cd44/ col4a1/ col4a2/ itgal* and *Icam1*.

Domain 10 was distinguished by an enrichment of mesenchymal populations—including pericytes, vascular smooth muscle cells (VSMCs), secondary crest myofibroblasts (SCMFs), and alveolar fibroblasts (AFs)—in the absence of detectable vascular structures (Fig. 6A, B). Interestingly, the cellular composition of this vascular-depleted domain is suggestive of a mesenchymal remodeling state that has been associated with pericyte-to-fibroblast transition in other tumor settings.(*19*) We performed CellChat v2, which incorporates spatial information to ensure cell interactions reflect biologically realistic physical distances. Our domain-restricted spatial CellChat v2 analysis suggested activation of stromal and matrix-associated communication programs, including PDGF and collagen signaling. This was supported by the spatial proximity of *Pdgfb*-and *Pdgfrb*-expressing cells and inferred pathway activities (Fig. 6D–F), and increased expression of the stromal marker *Sfrp1,* consistent with previous reports.(*19*) In addition, the presence of active collagen networks, along with HGF (Hepatocyte Growth Factor) and CD39 signaling pathways (Fig. 6D), suggests an ECM-rich microenvironment. These findings implicate Domain 10 as a potential contributor to tumor margin organization, although further validation with additional methodologies is needed.(*20, 21*)

In contrast, Domain 12 aligns with an immune-active niche, characterized by extensive immune cell infiltration and elevated expression of *Cxcr3, Cxcl9, Cxcl10*, and *Xcl1* (Fig. 6A, 6C). This chemokine signature indicates active recruitment of T cells to the tumor margin.(*22, 23*) Notably, PD-L1, TNF, and LCK signaling pathways were also enriched (Fig. 6D), supporting enhanced T cell–APC interactions and heightened immune checkpoint activity.(*24*) Collectively, these features suggest Domain 12 as a critical T cell-enriched regulatory micro-niche at the tumor boundary.(*23*)

Lastly, Domain 35 represents a myeloid-dominant niche enriched with pulmonary macrophages expressing *Gpnmb, Chil3*, and *Clec1b* (Fig. 6C), a signature characteristic of M2-like/lipid-associated macrophages (LAMs) involved in lipid metabolism and immunosuppression.(*25, 26*) This immunosuppressive function is further supported by active CD200 and TGFβ signaling (Fig. 6D).(*27, 28*) Notably, *Gpnmb-*expressing cells are in close spatial proximity to cells expressing integrins (*Spp1/Itga5/Itgb1*) and adhesion molecules (*Cd44/Col4/Icam1*) (Fig. 6G). This observation aligns with previous reports suggesting that *Gpnmb* may act as a dual ligand, engaging CD44 for survival signaling and interacting with integrins for myeloid-stromal adhesion.(*29, 30*) Taken together, these findings suggest Domain 35 as a structured peri-tumoral immunosuppressive myeloid micro-niche, although further validation through additional methodologies is needed.

### Tumor–adjacent–normal architecture is recapitulated in human lung cancer

Human lung cancer is substantially more complex than mouse models, largely due to its diverse genetic backgrounds and tumorigenic mechanisms. Consequently, results obtained from mouse studies often cannot be replicated in humans. To assess whether the spatial domain framework developed in the murine lung cancer model could be applied to human lung cancer, we implemented our analytic approach on a publicly available human lung cancer specimen (Fig. 7A).(*31, 32*) Consistent with our murine findings, BANKSY domains represented spatially coherent multicellular units that were well aligned with biologic structures. BANKSY clustering identified 30 spatial domains across the human lung cancer section. Spatial mapping revealed that these domains exhibited distinct, spatially restricted patterns corresponding to tumor-associated regions, preserved non-tumoral lung parenchyma, and discrete airway-or vessel-associated structures (Fig. 7A). Domain-specific highlight panels further confirmed that many domains consistently occupied distinct anatomical or tumor-associated locations.

**Fig. 7.**
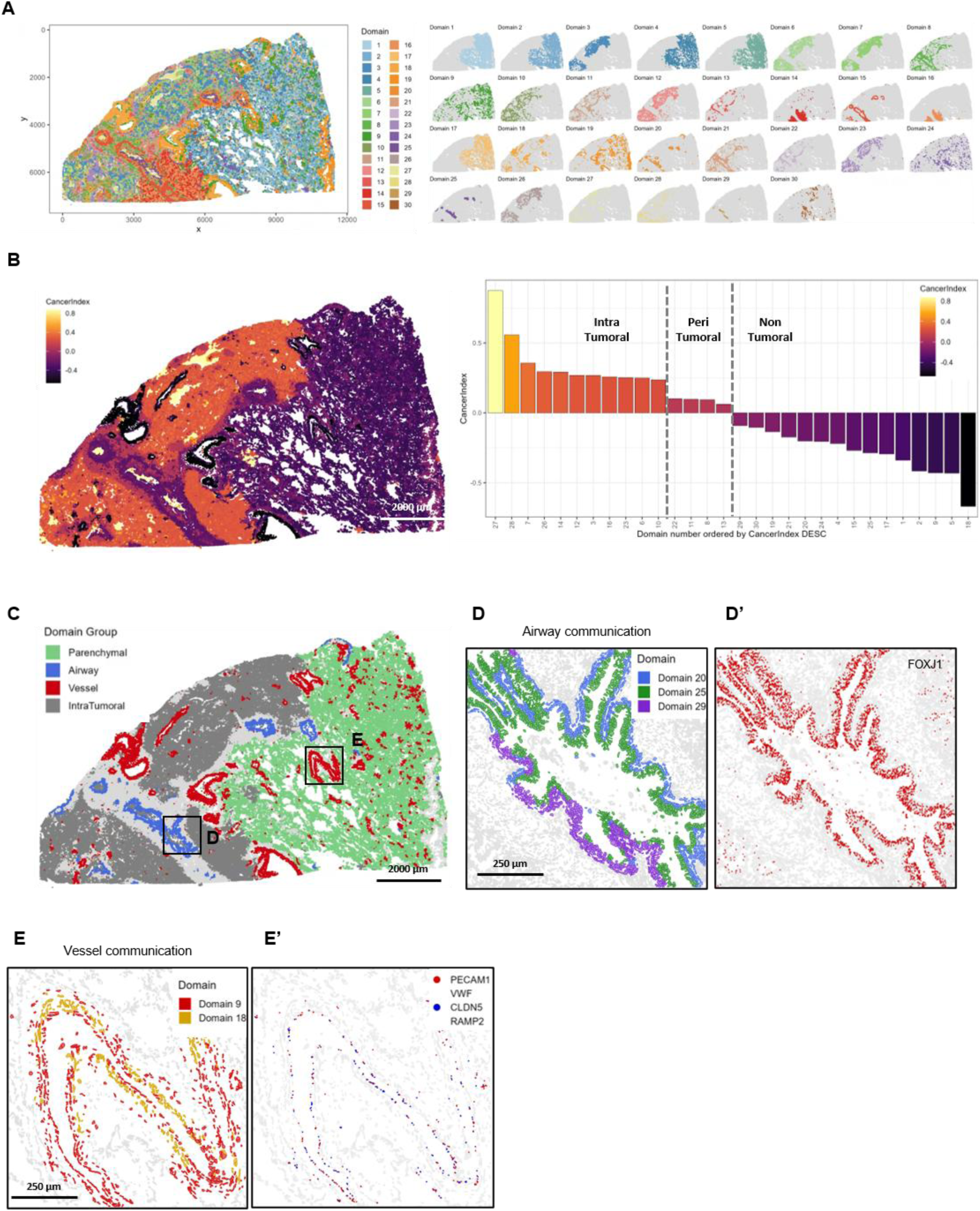
Human lung cancer spatial domain analysis validates the tumor ecosystem framework identified in the murine lung cancer model. **(A)** Spatial mapping of BANKSY-defined domains in a human lung cancer Xenium dataset. Thirty spatial domains were identified across the tissue section. The left panel shows the integrated domain map, and the right panels show individual domain highlight maps, demonstrating that domains localize to discrete tumor-associated or anatomical regions rather than being randomly distributed. **(B)** Spatial visualization of the domain-level Cancer Index in the human lung cancer specimen. Cancer Index values were calculated from ten selected hallmark programs and assigned to cells according to their corresponding BANKSY domain. Higher Cancer Index values mark tumor-associated regions, whereas lower values correspond to preserved non-tumoral tissue areas. Domain-level Cancer Index ranking. Human spatial domains were ordered by Cancer Index in descending order and classified into intra-tumoral, peri-tumoral, and non-tumoral groups. Domains with high Cancer Index values were assigned to the intra-tumoral compartment, domains with intermediate values to the peri-tumoral compartment, and domains with low values to the non-tumoral compartment. **(C)** Spatial map of domain group assignments in the human lung cancer section. **(D)** Non-tumoral domains were classified as parenchymal, airway-associated, or vessel-associated compartments, with intra-tumoral regions shown separately. **(D′)** Magnified airway-associated region showing Domains 20, 25, and 29 aligned with FOXJ1 expression. **(E)** Magnified vessel-associated region showing Domains 9 and 18 around vascular structures. **(E′)** Endothelial marker expression, including PECAM1, VWF, CLDN5, and RAMP2, corresponding to the vessel-associated Domains 9 and 18.

To calculate a domain-level Cancer Index, we refined the index to include conserved pathways shared with the mouse model as well as additional pathways that captured tumor-associated transcriptional variation specific to human lung cancer. For each domain, hallmark module scores were summarized to generate a Cancer Index as described in the Methods section. Spatial visualization of the Cancer Index showed marked regional variation across the human tissue section: higher scores concentrated in tumor-enriched regions, lower scores in preserved non-tumoral architecture, and intermediate scores in peritumoral areas (Fig. 7B). These findings demonstrate that Cancer Index-based stratification effectively distinguishes human spatial domains into intra-tumoral, peri-tumoral, and non-tumoral compartments, consistent with the results observed in the mouse model.

Next, we examined whether non-tumoral domains corresponded to distinct biologic lung structural compartments. Non-tumoral domains were further grouped into parenchymal, airway-associated, and vessel-associated compartments (Fig. 7C). Spatial mapping showed that parenchymal domains occupied broad regions of non-tumoral lung parenchyma, airway-associated domains were localized along airway-like structures, and vessel-associated domains formed focal or linear patterns around vascular structures. Notably, airway-associated regions exhibited a spatially ordered architecture that mirrored the organization of native airway structures (Fig. 7D). The airway-associated domains, including Domains 20, 25, and 29, showed layered organization along airway-like structures, which spatially corresponded to the expression of *FOXJ1*, a marker for ciliated airway epithelial cells (Fig. 7D′). Two domains, Domains 9 and 18, constituted the histologically matched vessel-associated region (Fig. 7E). These vessel-associated domains showed spatial correspondence with endothelial marker expression, including *PECAM1, VWF, CLDN5,* and *RAMP2* (Fig. 7E′). Altogether, these findings indicate that our spatial domain and Cancer Index analyses are applicable to human lung cancer tissue, though further validation with larger cohorts is warranted.

## Discussion

Domain-based analysis of spatial transcriptomics enables systematic dissection of the lung TME into recurrent, functionally specialized multicellular niches. By integrating high-plex Xenium spatial transcriptomics with spatially informed clustering, we identified distinct spatial domains that are transcriptionally coherent, topologically conserved, and reproducibly organized across both tumor-involved and non-tumoral lung sections. These results indicate that the lung TME is a spatially structured ecosystem governed by components of the local milieu that are variably dominant. (*33*),(*34*) Spatial domains constitute coordinated multicellular neighborhoods that capture recurrent patterns of local organization within the tumor microenvironment. (*35*),(*36*) Certainly, at crucial early moments in tumorigenesis or metastasis, there are likely instances of competing micro-domains overlapping incoherently, but in a given temporal snapshot, the vast majority of domains appear to be settled into stable local equilibria. This observation suggests that multicellular spatial niches may represent an important organizational layer of tumor ecosystems, complementing cell-centric views rather than replacing them. It is notable that the majority of discrete spatial domains resolved by this study could be readily characterized and identified, though a few failed to be elucidated, primarily due to their small number of composite cells. This comprehensive coverage highlights the suitability of spatial transcriptomics for systematic interrogation of cancer specimens. By simultaneously analyzing intra-tumoral, peri-tumoral, and non-tumoral regions within the same tissue context, our approach reveals spatial heterogeneity at both intratumoral and peri-tumoral levels that conventional bulk or dissociative methods cannot resolve.(*37*) This integrated strategy enables niche-specific biological interpretation and provides a unified framework for understanding how tumor-intrinsic programs interact with the surrounding structural and immune environments.

A key contribution of this study is the systematic characterization of both intratumoral and internodular heterogeneity, which appears to be associated with the organization of individual tumor nodules. By decomposing tumor nodules into their constituent spatial domains, we identified distinct architectural nodular subtypes characterized by unique micro-niche compositions, transcriptional programs, and signaling profiles (Fig. 4). These nodule subtypes likely reflect divergent evolutionary trajectories within a shared genetic background, consistent with prior observations of spatially constrained tumor evolution.(*38*),(*39*) Whether such diversity represents discrete evolutionary stages or adaptive responses to local microenvironmental pressure remains unresolved. Nonetheless, recognizing nodule-specific spatial architecture has significant translational implications. For instance, nodules enriched for inflammatory and immune-interacting niches may be more responsive to immune checkpoint blockade, while those dominated by stromal-or matrix components may require therapies targeting extracellular matrix remodeling or fibroblast activation. Nodules with high EMT signatures could prognosticate metastatic potential. Indeed, spatially encoded heterogeneity likely explains the limited efficacy of monotherapy, even in genetically homogeneous tumors, but also provides hope for more effective individualized combination therapies in the future.

Our analysis identified tumor-specific spatial domains that reside exclusively within intratumoral regions. Notably, we identified a tumor-associated mesothelial domain that forms a contiguous interface between tumor nodules and the pleural surface (Fig. 5). Although this domain shares lineage markers with non-tumoral normal mesothelium, it exhibits a distinct transcriptional profile enriched for inflammatory, stress-related, and extracellular matrix genes. These characteristics suggest that mesothelial cells adjacent to tumor tissue undergo spatially confined remodeling prior to the appearance of visible histological changes, potentially contributing to tumor progression and the formation of tumor boundaries.(*12*),(*40*) Notably, this process may be crucial to the development of malignant pleural effusions, which dramatically up-stage lung cancers and portend a grim prognosis.(*41*) In parallel, we identified an LEC–enriched domain localized at tumor margins, marked by robust CCR7–CCL21 signaling. This finding was further validated at the protein level by using IHC. CCR7 expression has been detected in migratory immune cells and subsets of malignant cells, implicating potential roles of this niche in both tumor antigen presentation and lymphatic tumor dissemination.(*42*) Notably, certain tumor nodule types exhibited a greater abundance of LEC-rich niches, indicating heterogeneous development of lymphatic micro-niches among nodules (Fig. 5F and 5G), which may influence intra-and peri-tumoral adaptive immune responses, as well as propensity for lymph node metastasis.(*42*),(*43*) Together, these findings highlight spatially defined, tumor-specific niches as critical interfaces with key roles in immune regulation and metastatic potential.

The peri-tumoral region emerges as a highly organized boundary, comprising multiple functionally distinct micro-niches rather than serving as a passive transition zone. We identified spatial domains enriched for stromal elements with activated adaptive immune cells (Domain 12) and immunosuppressive myeloid cells (Domain 35), strategically positioned along the tumor margin (Fig. 6). This spatial configuration implies adaptive compartmentalization in response to tumor expansion.(*23*),(*44*) Notably, GPNMB-positive macrophages are restricted to a peri-tumoral domain (Domain 35) characterized by robust immunosuppressive signaling.(*29*),(*45*) Thus, these macrophages appear to establish a spatially confined barrier enriched for adhesion and survival signals, potentially limiting immune cell infiltration into the tumor core (Fig. 6G). In contrast, adjacent peri-tumoral niches exhibit either immune-active chemokine expression or mesenchymal remodeling signatures (Domains 12 and 10, respectively, Fig. 6E and 6F), further highlighting the functional heterogeneity at the tumor boundary and illuminating potential roles for targeted immune-and chemotherapies(*23*),(*46*) Together, these findings support a model in which the peri-tumoral region serves as a structured interface integrating immune regulation, stromal remodeling, and tumor containment.(*44*) Future studies should further investigate intratumoral domains in relation to these peri-tumoral interfaces to determine how tumor-intrinsic states influence the formation and organization of boundary niches.

Importantly, our human lung cancer analysis demonstrates the feasibility of applying the spatial domain framework developed in the murine model can be extended to human disease. In the human Xenium dataset, Cancer Index–based domain classifications with BANKSY identified distinct spatial domains that paralleled our murine analysis, and organized the tissue section into intra-tumoral, peri-tumoral, and non-tumoral regions. (Fig. 7) In addition, it is worth noting that the low-Cancer Index non-tumoral domains recapitulated recognizable normal parenchymal, airway-associated, and vessel-associated structures, supporting the biological interpretability of spatial domains in human lung cancer tissue. Although human lung cancer is much more complex and diverse than mouse models, these findings support the broader concept that lung cancer can be understood as a spatial ecosystem comprising recurrent multicellular niches across species.

This study has several limitations. First, although our reanalysis of a single publicly available human lung cancer data supports the application of our spatial domain framework to human disease, larger human cohorts will be required to determine whether the same domain-level organization, Cancer Index-defined compartments, and structural micro-niches are conserved across patients, histologic subtypes, disease stages, and treatment contexts. Second, although reference-based cell-type annotation was rigorously validated, ambiguity in cellular classification remains an inherent challenge of spatial transcriptomics. Integrating matched single-cell RNA sequencing data in future studies will further refine cell-type resolution and strengthen mechanistic interpretation.

Despite these limitations, our study presents a conceptual and analytical framework for interpreting the lung tumor microenvironment as an organized spatial ecosystem composed of recurrent multicellular niches. Utilizing high-resolution spatial transcriptomics integrated with spatially informed domain analysis, we show that tumor heterogeneity and behavior are fundamentally shaped by the organization of spatial domains and their assembly into tumor nodules. This framework connects canonical lung architecture, tumor-specific micro-niches, peri-tumoral boundary states, and human validation data into a cohesive model of lung cancer spatial organization, with important implications for immune responsiveness, tumor progression, and therapeutic vulnerability. Taken together, these results provide a foundation for integrating spatial ecosystem states into tumor classification frameworks, ultimately advancing precision oncology.

## Materials and Methods

### Study Design

The objective of this exploratory study was to determine whether spatially resolved multicellular domains could define higher-order organization within the lung tumor microenvironment. Three independently tumor-bearing mice served as biological replicates, and one tissue section from each mouse was analyzed. No formal power calculation was performed because the study was designed as a discovery-oriented spatial transcriptomic analysis. Sample size was not altered after data collection began. Cell-and domain-level inclusion and exclusion criteria were established before downstream comparative analyses and are described below. Investigators were not blinded during computational analysis. The publicly available human lung cancer specimen was used as an external exploratory assessment of cross-species applicability rather than as an independent cohort-level validation.

### Mouse Models and Tumor Induction

Primary lung adenocarcinomas were induced in Scgb1a1-CreERT2;LSL-Kras^G12D^;Trp53 ^flox/flox^ (KP) mice by tamoxifen administration, and tumor cells were isolated and established as previously described.(*5*) Established KP lung cancer cells were transduced with a lentiviral vector encoding a zsGreen-minOVA fusion cassette, followed by clonal expansion to ensure uniform expression. These engineered cells were orthotopically transplanted into syngeneic recipients via intratracheal delivery. Four weeks post-transplantation, lungs were harvested, formalin-fixed, and paraffin-embedded (FFPE). Three individual mice were analyzed as biological replicates. One representative FFPE lung tissue section was selected from each mouse and designated as samples 4w1, 4w2, and 4w3, respectively. Serial sections were stained with H&E to identify regions of interest (ROIs) encompassing both tumor nodules and adjacent non-malignant parenchyma. Sections immediately adjacent to H&E-validated ROIs were then mounted on Xenium slides for downstream spatial transcriptomic Analysis.

### Xenium in Situ Spatial Transcriptomics

Xenium in situ spatial transcriptomic profiling was performed at Northwestern University’s NUSeq Core Facility using the Xenium platform (10x Genomics) with Xenium Prime chemistry. Briefly, 5-µm FFPE tissue sections were mounted onto Xenium slide capture areas and incubated at 42 °C for 3 hours, followed by overnight drying. Slides were stored at room temperature in a desiccation chamber prior to processing. Slide preparation and Xenium Analyzer loading were carried out using the Xenium Slide and Sample Preparation Reagent Kit (PN1000460), Xenium Decoding Consumables (PN-1000487), and Xenium Decoding Reagents (PN-1000461), strictly according to the manufacturer’s instructions.

Slides were first deparaffinized and de-crosslinked following protocol CG000580. Xenium in situ gene expression was then performed using a panel of probes targeting a total of 5,054 genes, comprising 5,006 pre-designed genes from the 10x Genomics catalog and 48 custom-selected targets selected to represent epithelial, immune, stromal, and endothelial lineages. Probe hybridization was performed overnight, followed by probe ligation and enzymatic signal amplification according to manufacturer protocols (CG000582, CG000749, and CG000760). Autofluorescence quenching and nuclear staining were performed prior to imaging.

Slides were imaged on the Xenium Analyzer (instrument software version 3.2.1.2), and raw fluorescence images were processed using the instrument’s onboard primary analysis pipeline to decode transcript identities and assign spatial coordinates to individual RNA molecules. Cell and nucleus segmentation were performed using the 10x Genomics multimodal segmentation algorithm within the Xenium Onboard Analysis (XOA) pipeline version xenium-3.2.0.7 (10x Genomics; https://www.10xgenomics.com/support/software/xenium-onboard-analysis/latest). This proprietary deep learning-based framework integrates membrane boundary staining, interior RNA staining (18S rRNA), and DAPI nuclear staining to delineate cellular compartments hierarchically. Membrane boundary staining was prioritized when detected; otherwise, whole-cell boundaries were defined using a deep learning-assisted expansion of DAPI-identified nuclei to the interior stain edge. Across all tissue sections, more than 90% of cells were segmented using the nuclear-to-interior stain edge strategy, whereas only a small minority (∼1–2%) required predefined nuclear expansion. Following segmentation, spatially decoded RNA transcripts were assigned to segmented cells to generate nuclear and whole-cell polygon boundaries, along with corresponding cell-by-gene count matrices, for downstream analysis. In total, the XOA pipeline yielded 110,436 cells for sample 4w1, 168,368 cells for sample 4w2, and 279,720 cells for sample 4w3.

### Xenium Data Processing, Normalization, and Differential Testing

To systematically decode the spatial ecosystem of the lung TME, we implemented a high-plex in situ transcriptomic analysis using a rigorous quality control and integration framework following XOA pipeline processing, with data visualization performed using custom plotting functions in R (v4.5.1). To preserve biological signal integrity while minimizing technical artifacts, cells were filtered using segmentation-and coverage-based criteria, including cell segmentation, negative-control–informed library metrics, and minimum transcript and detected gene thresholds. (*47*)

To address potential technical limitations of the Xenium Prime 5K platform, such as transcript diffusion and optical bleed-through, Xenium-native robustness analyses were conducted in place of droplet-based background correction tools. Specifically, a Baysor parameter sweep was performed to optimize the min_molecules_per_cell threshold. By directly comparing the original Xenium segmentation matrix with the Baysor resegmentation results, the analysis confirmed that the identified spatial domains reflect genuine biological structures rather than artifacts driven by low-quality cells, extracellular RNA background, or chimeric transcript assignments.

XOA pipeline output was imported into Seurat (v5.3.1) function *LoadXenium*, retaining molecule-level coordinates (Qv ≥ 20), cell centroids, and both cell-and nucleus-level segmentation boundaries, which were consolidated into a single Seurat object per tissue section. Per-cell segmentation metrics derived from XOA pipeline output tables (including cell area, nucleus area, and nucleus count) were appended to the Seurat metadata. Additional quality control metrics were computed upon data transfer to a SpatialFeatureExperiment (v1.12.1) object using the scuttle package (v1.20.0) function *addPerCellQCMetrics*, which quantified total counts, detected features, and the proportion of counts attributable to negative control probes, negative control codewords, and unassigned codewords.(*48–50*) Cells exceeding a 2.5% threshold for combined negative control signal were excluded. Library-size outliers were identified using median absolute deviation–based filtering on log-transformed per-cell counts (nmads = 3, lower tail), and spatially isolated cells were removed using a k-nearest neighbor distance criterion (k = 5), retaining only cells within the main tissue region (max neighbor distance ≤ 200 and min neighbor distance ≤ 50 in spatial coordinate units). In Seurat, additional filtering excluded cells with abnormal nuclear segmentation, and analyses were restricted to cells that exhibited a single nucleus (nucleus count = 1) to minimize artifacts arising from segmentation irregularities or overlapping nuclei. Following filtering, transcript and gene counts were normalized to cell and nuclear area to derive count-density metrics, which were visualized to assess distribution patterns prior to merging retained cells (4w1 = 107,862; 4w2 = 163,522; 4w3 = 265,598) for downstream normalization and integration.

After sample-level quality control, the three QC-filtered Seurat objects were subsequently merged while preserving sample identity and spatial fields of view, yielding a combined dataset of 536,982 cells across 4w1, 4w2, and 4w3. Post-merge quality assessment confirmed retention of single-nucleus cells (nucleus count = 1 for all cells) and preserved distributions of transcript counts, detected genes, and segmentation metrics across samples. Cells in which the nucleus area equaled the total cell area (nucleus-to-cell ratio = 1), suggestive of segmentation edge artifacts, were identified and removed, resulting in a final merged dataset of 535,796 high-quality cells for downstream normalization and integration. Distributions of transcript counts, feature counts, and morphology-derived covariates were re-evaluated on the merged object to ensure consistency across sections prior to integrated analysis.

Given that cell-level quality control and normalization were completed prior to downstream modeling, we next considered whether additional gene-level filtering was warranted. Xenium utilizes a curated, probe-based gene panel; therefore, no additional gene-level filtering based on detection frequency or minimum count thresholds was applied. Quality control focused on removal of control probes and low-quality cells, and all 5,054 panel genes were retained for downstream analyses to preserve cell-type-restricted and spatially localized transcripts. Gene expression counts in the Xenium assay were normalized in Seurat using library size–based log normalization (*NormalizeData* function; method = “LogNormalize”), with the scale factor set to the median per-cell transcript count across the merged dataset (scale.factor = median(nCount)), thereby rescaling each cell to a dataset-specific Xenium library. For variance stabilization and downstream dimensionality reduction, the merged object was subsequently processed using SCTransform (vst.flavor = “v2”; ncells = 5000; return.only.var.genes = FALSE; do.correct.umi = TRUE; do.center = TRUE; do.scale = FALSE), generating an SCT assay that was used for principal component analysis, Uniform Manifold Approximation and Projection (UMAP) embedding, and differential expression analysis between cell type clusters and domains. Notably, prior to differential expression testing, the *PrepSCTFindMarkers* function was applied to ensure consistent UMI model parameters across SCT layers in the merged object, as recommended for multi-sample SCT workflows. Differential expression was performed using Seurat’s *FindAllMarkers* function (assay = “SCT” or “BANKSY” when applicable, slot = “data”, test.use = “wilcox”; min.pct = 0.2; logfc.threshold = 0.25; min.cells.group = 20), and p-values were adjusted for multiple comparisons using the Benjamini–Hochberg and considered significant at a false discovery rate < 10%.(*48, 51, 52*)

### Human lung cancer Xenium dataset analysis

Publicly available Xenium human lung cancer data were obtained from the 10x Genomics dataset “Xenium v1 and Xenium Prime 5K for FFPE Human Lung Cancer.” The dataset was generated from an FFPE human lung cancer tissue section using the Xenium platform and includes spatial gene expression data generated with the Xenium Human Lung Gene Expression Panel.(*32*)

Processed Xenium output files were downloaded from the 10x Genomics public data portal and imported into Seurat using the LoadXenium function. The human dataset was analyzed using the same spatial domain analysis framework applied to the mouse Xenium dataset, where applicable, including normalization, preservation of spatial coordinates, BANKSY-based spatial domain detection, and Cancer Index scoring. Human non-tumoral domains were further annotated into parenchymal, airway-associated, and vessel-associated compartments based on spatial localization and marker gene expression.

### Cell-Type Classification

Cell type annotation was performed using a reference-based strategy implemented in SingleR (v2.12.0).(*53*) As an external reference, we utilized the LungMAP Mouse Lung CellRef v1.1, a curated single-cell atlas of murine lung comprising 71,439 cells and 26,727 genes with hierarchical annotations spanning lineage-and cell type– specific levels.(*7*) Both our Xenium dataset and the reference were converted to a SingleCellExperiment object and the LungMap reference was used to annotate the Xenium dataset at “cell-type level1” resolution, which includes 31 pulmonary cell types representing epithelial, endothelial, immune, and mesenchymal compartments. Only genes shared between the reference and Xenium panel were retained for classification (n = 4,784 intersecting genes). The *SingleR* function was run using log-normalized expression values (assay.type.test = “logcounts”; assay.type.ref = “logcounts”) with de.method = “wilcox”, as recommended for single-cell reference datasets. This approach assigns cell type labels to individual cells by comparing each Xenium profile to reference-derived marker signatures. The resulting annotations encompassed 31 LungMAP-defined pulmonary cell types, including epithelial populations (AEC, AT1/AT2, AT2, Basal, Ciliated/Deuterosomal, Secretory, PNEC, Sox9 Epi), endothelial populations (CAP1/EPC, CAP2, VEC, LEC), immune populations (AM, IM, Monocyte, Neutrophil, NK, B, T, ILC, DC, Mast, Basophil, Megakaryocyte/Platelet), and mesenchymal populations (AF1, AF2, ASMC, Pericyte, SCMF, Mesothelial, VSMC, Chondrocyte).

To resolve transcriptional heterogeneity within cell types of interest, we performed sub-clustering of immune and mesenchymal cell populations defined at CellType Level 1. For each selected lineage, raw counts were re-normalized using *SCTransform*, followed by dimensional reduction and unsupervised clustering across all three tissue sections combined. This re-normalization step was implemented to optimize detection of within-cell-type biological variation that may not be fully captured by global normalization performed on the complete dataset. Cluster resolution was selected empirically for each cell-type based on the presence of robust and reproducible differential gene expression signatures, ensuring that resulting subclusters were supported by sufficient marker genes to define biologically meaningful states (the majority of sub-clusters had at least 10 differentially overexpressed genes.

### Spatial Domain Identification using BANKSY

To define spatial domains within the tumor microenvironment, we applied the BANKSY algorithm (v1.6.0) to the merged, QC-filtered dataset using the Seurat wrapper function *RunBanksy*. BANKSY was run on log-normalized expression values from the Xenium assay (performed utilizing Seurat’s *LogNormalize* function and scaled to the median per-cell transcript count, as described above). (*6*) Spatial feature construction incorporated cell centroid coordinates (x-centroid, y-centroid) and a geometric neighborhood defined by k-nearest neighbors (k = 18), with spatial smoothing parameter λ = 0.8 extracted on two spatial dimensions. Multi-sample processing was performed jointly across all three tissue sections using group = orig.ident and split scaling to preserve sample-specific structure while enabling integrated domain detection.

Dimensionality reduction was performed using PCA on the BANKSY assay (20 components), followed by UMAP embedding for visualization. Shared nearest-neighbor graph construction and community detection were then carried out using Seurat’s *FindNeighbors* and *FindClusters* functions across a range of resolutions. A resolution sweep was conducted from 1.0 to 2.0, and a resolution of 1.6 (BANKSY_snn_res.1.6) was selected based on cluster stability and biological interpretability, yielding 46 distinct spatial domains representing multicellular functional units. The robustness of these clustering parameters was validated using the Adjusted Rand Index (ARI) and alluvial analyses across neighboring resolutions. These domains represent multicellular functional units rather than single cell types and were subsequently evaluated for cell type composition and cross-sample consistency. To ensure reproducibility, domain labels were treated as numeric identifiers and explicitly encoded as factor variables to maintain consistent downstream ordering.

### Cancer Index-Scoring and Domain Annotation

To stratify spatial domains into biologically interpretable tumor-associated compartments, we developed a Cancer Index-scoring framework based on the transcriptional activity of hallmark cancer-related programs. For the mouse lung cancer dataset, spatial domains were scored using selected MSigDB Hallmark gene sets, including epithelial–mesenchymal transition, hypoxia, KRAS signaling down, MYC targets V1/V2, apoptosis, TNFA signaling via NFκB, IL6–JAK–STAT3 signaling, and glycolysis.(*8*) For the human lung cancer dataset, Cancer Index-scoring was performed using a fixed set of ten Hallmark marker sets selected for domain-level annotation: unfolded protein response, p53 pathway, complement, TNFA signaling via NFκB, coagulation, KRAS signaling down, apical junction, Notch signaling, IL6–JAK–STAT3 signaling, and mitotic spindle.(*8*)

For each dataset, enrichment scores for the selected Hallmark gene sets were calculated for each spatial domain and standardized as z-scores across domains. Scores were directionally aligned where necessary so that higher values consistently reflected stronger tumor-associated transcriptional activity. The standardized pathway scores were then averaged with equal weighting to generate a composite Cancer Index for each domain. Based on the distribution of Cancer Index values, spatial localization, and histologic context, spatial domains were classified into ecosystem categories, including intra-tumoral, peri-tumoral, and non-tumoral compartments. In the human lung cancer dataset, non-tumoral domains were further annotated as parenchymal, airway-associated, or vessel-associated compartments according to their spatial organization and marker gene expression patterns. The spatial accuracy of the constructed Cancer Index was quantitatively validated through a pixel-level concordance analysis against ground-truth tumor regions defined by HALO (as described in the next section). When translating this index to human datasets, the gene-set composition was modified to include detectable and informative Hallmark pathways that account for species-specific differences, ortholog mapping, and platform-dependent gene coverage.

### Post-Xenium H&E Staining and HALO Image Analysis

Following Xenium imaging, the same tissue sections were processed for post-run H&E staining according to the manufacturer’s protocol, scanned on an Evident SlideView VS200 at 20× magnification, and saved to .vsi format. The images were converted to ome.tif at the original pixel size and exported from QuPath (v0.7.0) without downsampling. These ome.tif images were imported into the HALO Image Analysis platform (v4.2.6399.280; Indica Labs), regions of interest (ROIs) were drawn around tissue, and negative regions were created around artifacts and non-target tissue. Subsequently, a two-step tissue classifier pipeline was created. In step 1, a tissue/no-tissue random forest classifier was trained to exclude non-tissue areas. The tissue class was then input for the second step, a dense net deep learning classifier, which was trained to identify tumor and non-tumor lung tissue. To create this classifier, 66 training regions across 9 slides from individual mice in our mouse model (including the current Xenium tissue sections) were used.

Image registration was performed using STalign (v1.0) in a Python 3.10.20 environment. Xenium morphology images served as the fixed reference image, whereas H&E images served as the moving image. Landmark-guided affine registration was performed using manually annotated corresponding tissue landmarks (n=10) to align the H&E image to the Xenium DAPI image. The resulting affine transformation matrix was applied to all HALO segmentation objects, thereby transforming pathology-derived tumor annotations into Xenium image space. All subsequent tumor-mask generation, rasterization, visualization, and concordance analyses were performed on the full-resolution Xenium image grid.

To quantify concordance between pathology-derived and transcriptomics-derived tumor annotations, we compared tumor masks generated from HALO-based segmentation and Xenium Banksy domain–Cancer Index classification on full-resolution Xenium DAPI images. HALO tumor annotations were exported as object-level segmentation masks from H&E images and spatially aligned to the Xenium coordinate system using the affine transformation derived from image registration. Tumor annotations from both HALO and Xenium analyses were represented as polygon-based segmentations, converted to Xenium DAPI pixel coordinates using the native Xenium image resolution (0.2125 µm/pixel), and rasterized into binary masks within Xenium image space using the rasterio package in Python.

Following mask generation, HALO and Xenium tumor masks were represented on the same full-resolution pixel grid and compared on a pixel-by-pixel basis, and tumor burden was calculated from the total number of tumor pixels and converted to physical area using the native Xenium pixel area (0.2125² µm² per pixel). A confusion matrix was generated to quantify the agreement among shared tumor pixels, HALO-only tumor pixels, Xenium-only tumor pixels, and background. For quality control and visualization, HALO and Xenium tumor masks were overlaid on the Xenium DAPI image to assess the spatial distribution of concordant and discordant tumor regions. From the confusion matrix, balanced accuracy and the Matthews correlation coefficient (MCC) were calculated to assess overall agreement between pathology-derived and transcriptomics-derived tumor annotations. Spatial concordance was quantified using the Dice coefficient, and tumor burden concordance was evaluated by calculating total tumor area for each method, shared tumor area, and the ratio of Xenium-derived to HALO-derived tumor area.

### Spatial Community Analysis

We characterized the architecture of the lung ecosystem by defining “structural communities” and “tumor nodule types” based on spatial domain composition. For non-tumoral regions, we grouped domains into anatomical structures (e.g., BALT, airway, vessel) by quantifying the enrichment of lineage-specific cell types and canonical marker expression within each domain. Conversely, we deconstructed intratumoral heterogeneity by classifying tumor nodules into five distinct pattern types (Types 1–5). This classification was driven by the dominant prevalence of specific intra-tumoral spatial domains (e.g., Domain 8 for Type 1; Domain 7 for Type 2), revealing that tumor nodules are not uniform but are organized into distinct micro-niches with unique stromal and immune compositions. Pairwise dissimilarities in intra-tumoral domain composition between these nodules were measured using the Bray-Curtis distance metric. To rigorously evaluate the statistical separation of these nodule types, Permutational Multivariate Analysis of Variance (PERMANOVA) and Principal Coordinate Analysis (PCoA) were performed, demonstrating that the nodule archetype accounted for a significant majority of the variance in spatial domain composition even after adjusting for sample identity.

### Cell-Cell Communication Analysis

To decode the signaling logic governing these micro-niches, we inferred ligand-receptor interactions using CellChat v2 in spatial mode.(*54*) We parameterized spatial constraints directly from tissue coordinates, computing the 1-nearest neighbor (k=1) distance distribution within each domain. We set the interaction range (‘contact.rang’) to the domain-specific third quartile (Q3) distance, with a scaling factor of 1.05 (‘scale.distanc’) to robustly capture local signaling neighborhoods. Communication probabilities were estimated using the ‘truncatedMean’ method with a trim parameter of 0.1. We filtered interactions based on a minimum of 10 cells per interacting group and aggregated significant signaling pathways (p < 0.05) to construct domain-specific communication networks.(*54*)

### Differential Expression and Pathway Analysis

We elucidated the molecular mechanisms driving domain identity by performing domain-specific differential expression analysis. We utilized the ‘FindAllMarkers’ function in Seurat, identifying genes significantly upregulated in specific domains compared to the rest of the tissue (log2 fold change > 0.25, FDR < 0.05). To biologically contextualize these signatures, we conducted domain-specific intercellular communication analysis using CellChat v2(*54*), inferring spatially constrained ligand–receptor interactions within each BANKSY-defined domain and summarizing signaling activities at the pathway level.

### Visualization and Statistical Analysis

We visualized the high-dimensional landscape of the tumor ecosystem using Uniform Manifold Approximation and Projection (UMAP) on the BANKSY-integrated feature space. Spatial mapping of domains and gene expression was performed using custom plotting functions in R. Statistical comparisons of domain abundance and gene expression between groups were assessed using the Wilcoxon test, with p-values adjusted for multiple hypothesis testing using the Benjamini-Hochberg method. Correlations between spatial features were evaluated using Pearson’s correlation coefficient.

## Data and Code Availability

The raw and processed Xenium spatial transcriptomics data generated in this study have been deposited in the Gene Expression Omnibus (GEO) under accession number GSE319943. The complete computational analysis pipeline, including scripts for BANKSY clustering, Cancer Index-scoring, and CellChat inference, is available on GitHub at (https://github.com/Minhyung83/Xenium_Spatial_data_Lung_Cancer).

## Acknowledgements

The authors acknowledge the Research Tissue Imaging Core at the University of Illinois at Chicago for providing imaging services.

## Author contributions

B.B., S.K., and K.K. performed the mouse experiments, analyzed data, and drafted the manuscript. M.K. and S.Y. conducted bioinformatic analyses and contributed to drafting the manuscript. J.E. contributed to the discussion and edited the manuscript. C.A. and G.P. conceptualized the study and supervised the data analysis. All authors edited the manuscript and provided feedback.

## Data and materials availability

All data generated or analyzed during this study are included in this published article and its supplementary information files.

## Ethics approval and consent to participate

All mouse strains were bred in a specific pathogen-free facility maintained by the University of Illinois at Chicago. All mouse experiments were approved by the Institute Animal Care and Use Committees (IACUC) of the University of Illinois at Chicago and Jesse Brown VA Medical Center.

*This manuscript was prepared with minimal assistance from ChatGPT to ensure its integrity. The authors thoroughly reviewed and validated the content.

## Supplementary Figure legends

**Fig. S1.**
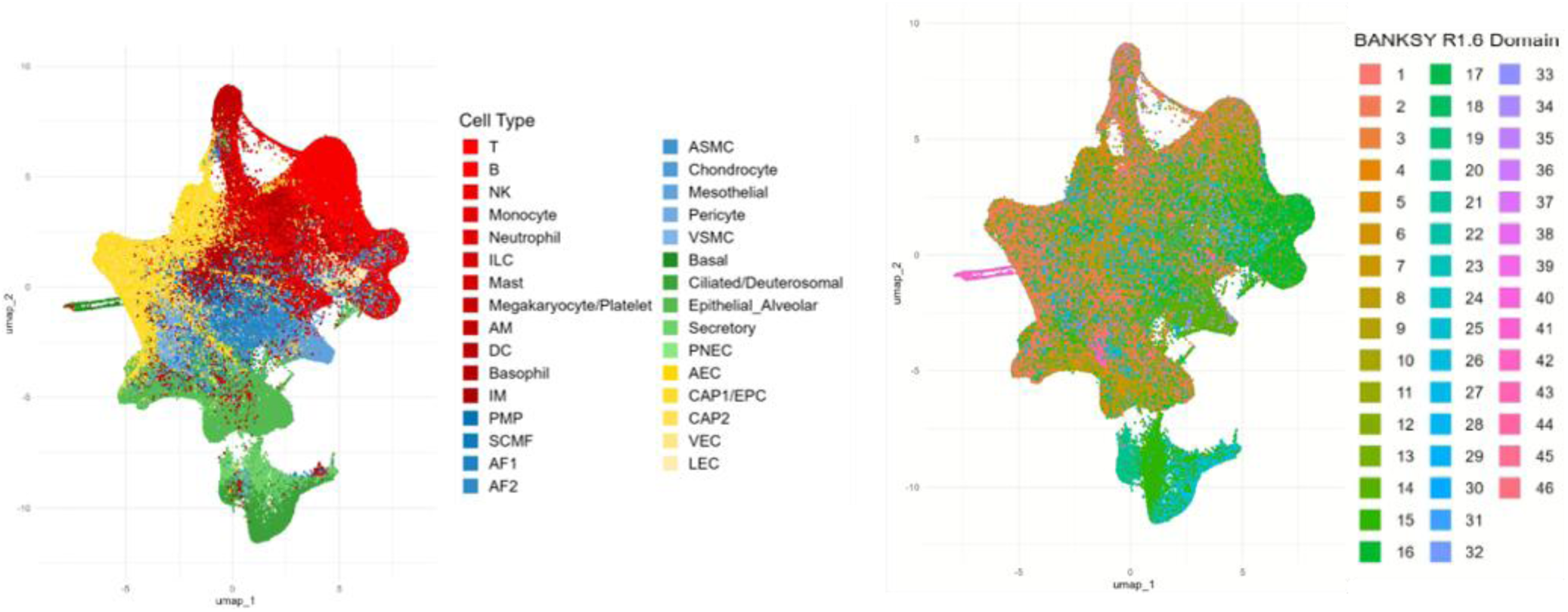
UMAP visualization of cell-type annotations and BANKSY-derived spatial domains. (A) UMAP embedding of Xenium cells colored by reference-based cell-type annotations. Major epithelial, endothelial, immune, and mesenchymal cell populations are shown. (B) UMAP embedding of the same cells colored by BANKSY spatial domain assignments at resolution 1.6. The broad distribution of domains across transcriptional space indicates that BANKSY domains represent multicellular spatial units rather than conventional single-cell-type clusters.

**Fig. S2.**
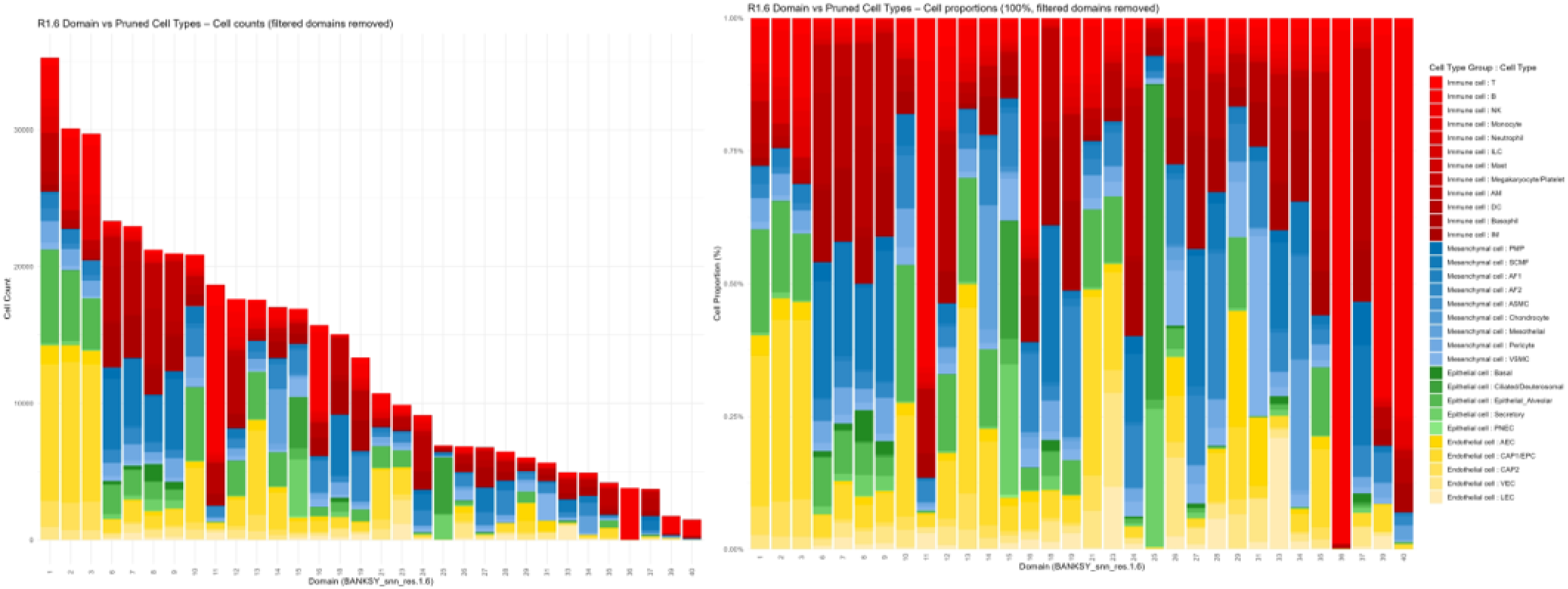
Cell-type composition of BANKSY-derived spatial domains. (A) Stacked bar plot showing the number of cells assigned to each BANKSY spatial domain at resolution 1.6, colored by reference-based cell-type annotation. (B) Relative cell-type composition of each BANKSY spatial domain, shown as normalized proportions. Together, these plots demonstrate that spatial domains are composed of distinct mixtures of epithelial, endothelial, immune, and mesenchymal cell populations rather than single cell types.

**Fig. S3.**
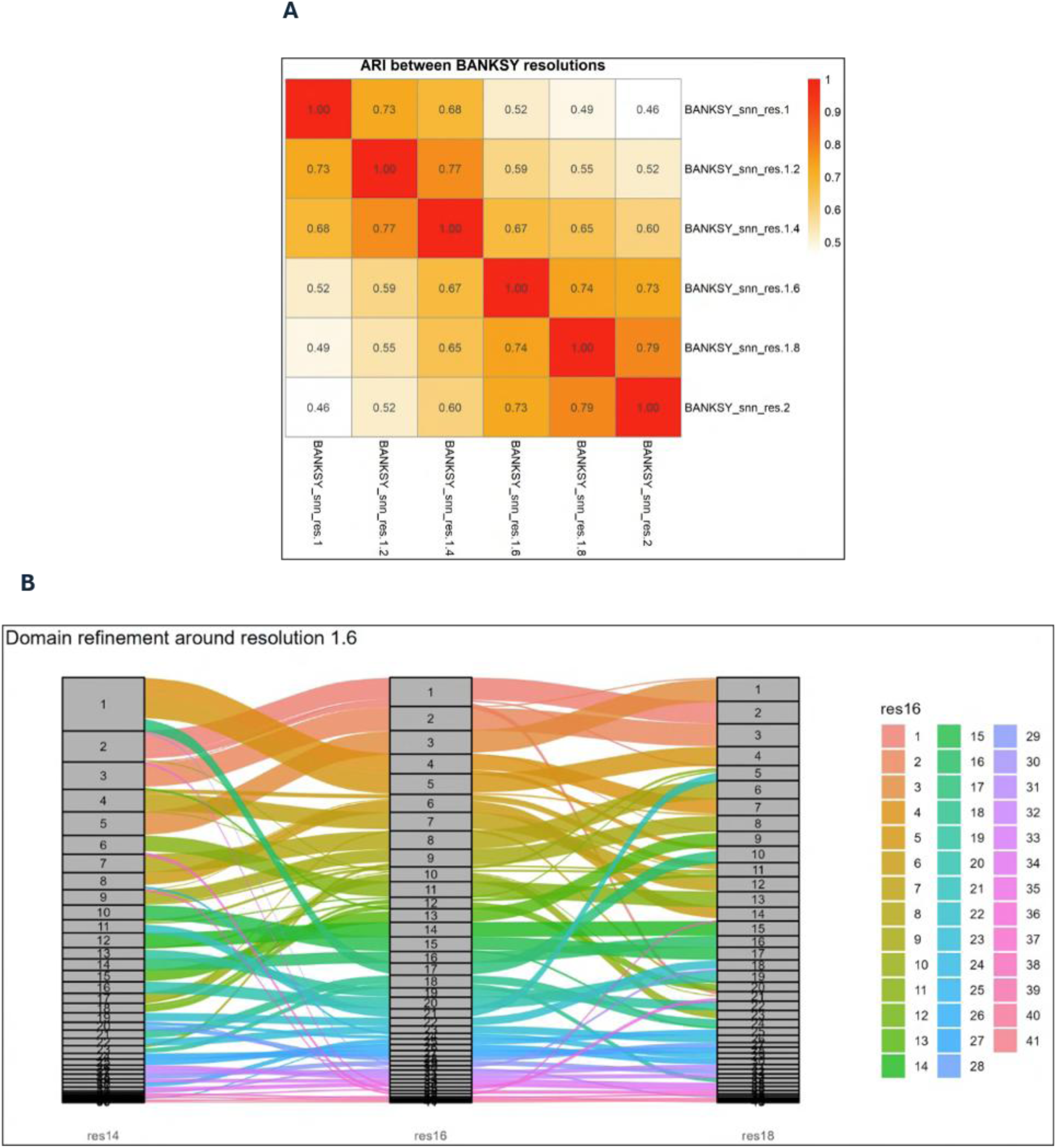
Robustness analysis of BANKSY spatial domain resolution. (A) Pairwise adjusted Rand index (ARI) between BANKSY clustering results generated across different resolution values. Higher ARI values between neighboring resolutions indicate stable domain assignments across parameter settings. (B) Alluvial plot showing domain relationships across resolutions 1.4, 1.6, and 1.8. Domains at the selected resolution 1.6 were largely preserved across neighboring resolutions, with higher resolution mainly producing hierarchical refinement of existing domains.

**Fig. S4.**
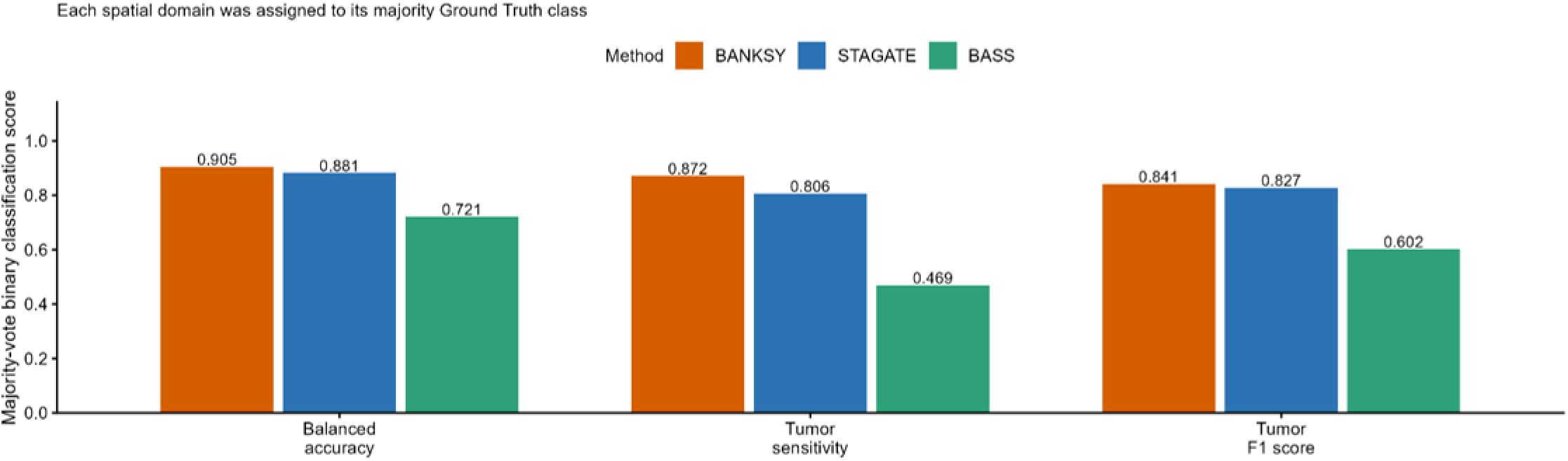
Recovery of Ground Truth tumor Structure by alternative spatial clustering methods. Bar plots comparing the performance of BANKSY, STAGATE, and BASS for tumor-region classification. Each spatial domain was assigned to its majority ground-truth class, and classification performance was evaluated using balanced accuracy, tumor sensitivity, and tumor F1 score. BANKSY achieved the highest performance across all three metrics, supporting its selection for downstream spatial domain analysis.

**Fig. S5.**
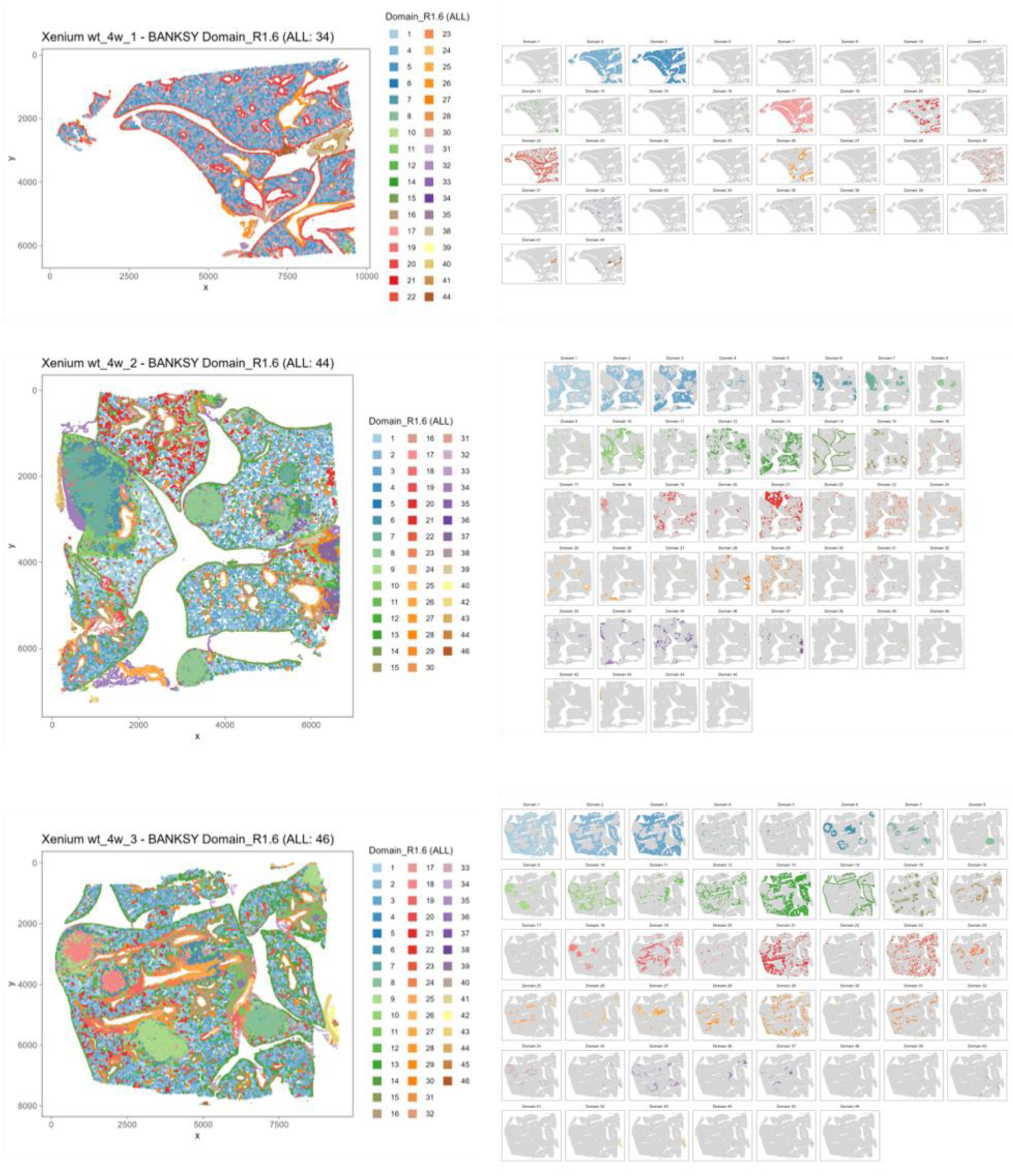
Sample-wise spatial distribution of BANKSY-derived domains. (A–C) Spatial maps of BANKSY domain assignments at resolution 1.6 for individual Xenium lung tumor sections 4w1, 4w2, and 4w3, respectively. (D–F) Domain-specific spatial maps showing the localization of each BANKSY domain within the corresponding tissue section. These plots demonstrate that major spatial domain structures are reproducibly detected across independent tumor-bearing lung samples while preserving sample-specific tissue architecture.

**Fig. S6.**
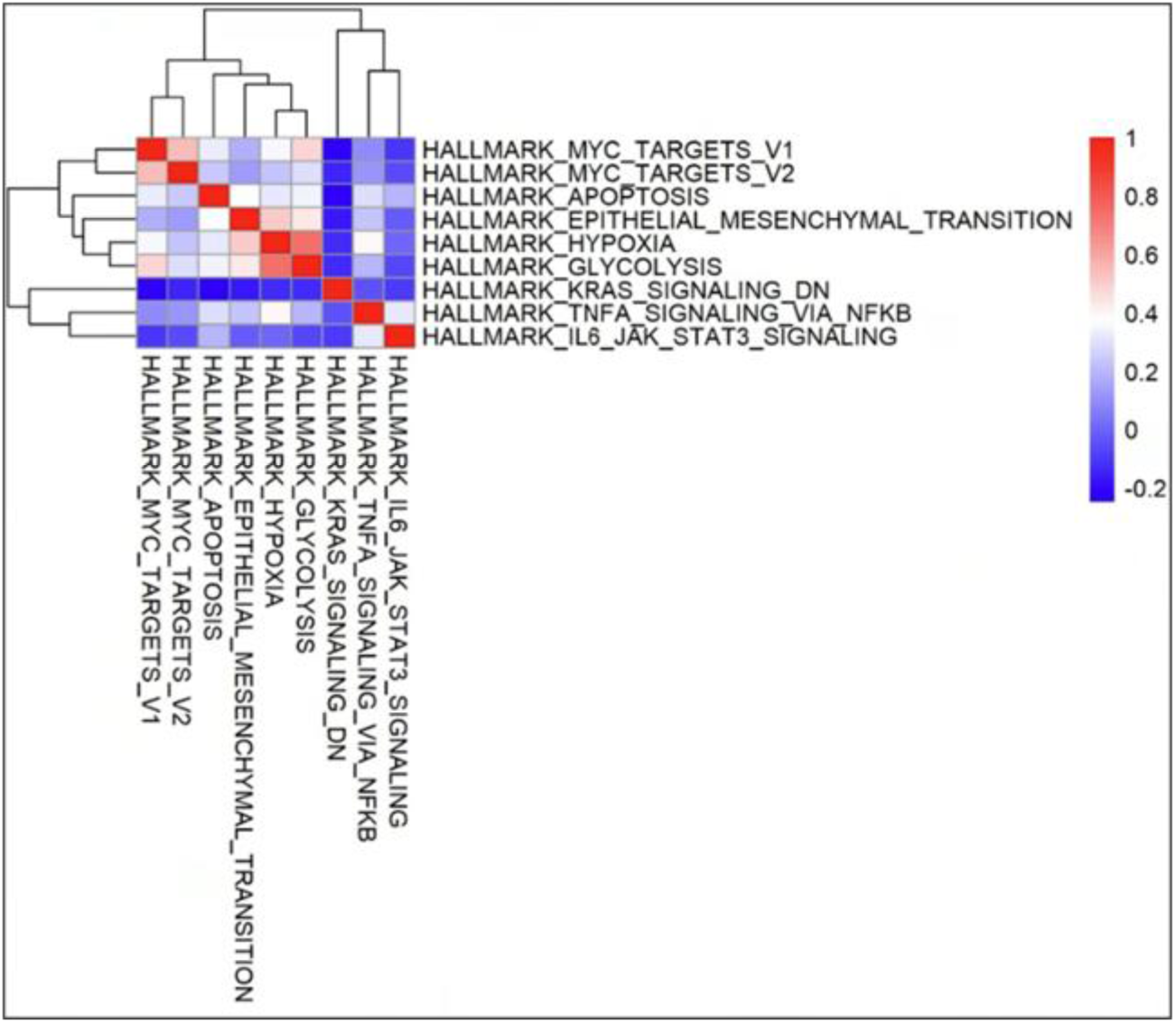
Correlation analysis of Hallmark pathway components used for CancerIndex construction. Clustered correlation heatmap showing pairwise correlations among Hallmark pathway activity scores included in the mouse CancerIndex. Partial correlations among pathway scores indicate that the CancerIndex integrates multiple, non-redundant tumor-associated programs rather than being driven by a single Hallmark pathway.

**Fig. S7.**
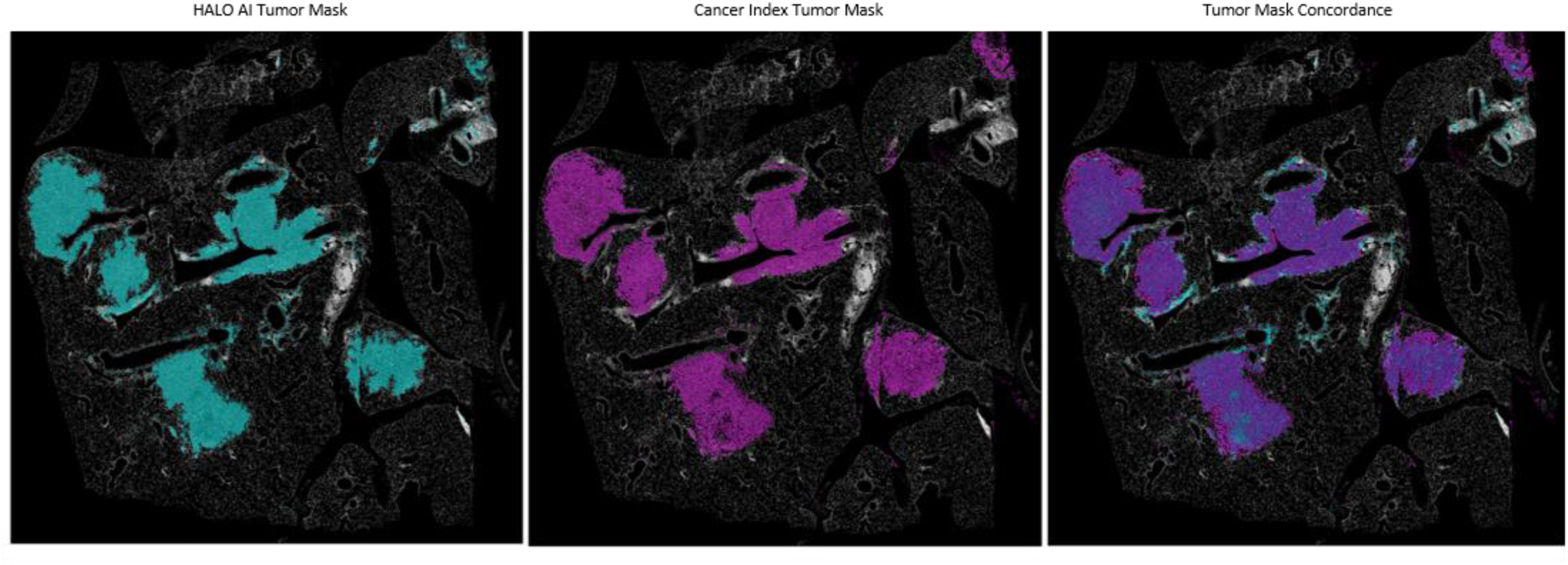
Validation of BANKSY-based tumor-domain classification against HALO-derived ground-truth tumor annotations in sample 4w3. Representative spatial comparison of HALO-derived ground-truth tumor regions, BANKSY/Cancer Index–derived tumor classification, and their merged overlay in the 4w3 tissue section. This analysis supports the reproducibility of the spatial concordance shown in Fig. 2D, which presents the corresponding 4w2 validation.

---

**Table S1.**
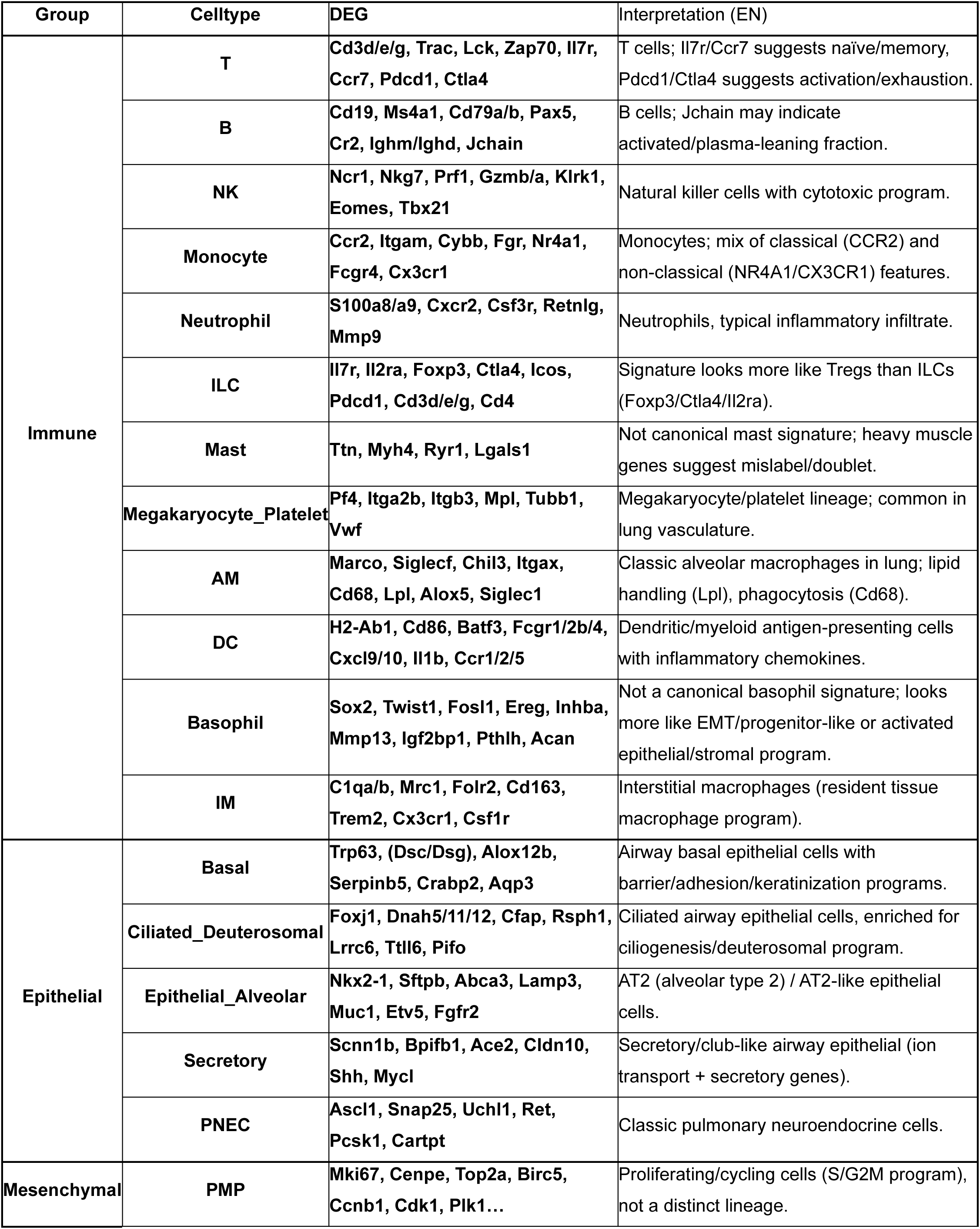
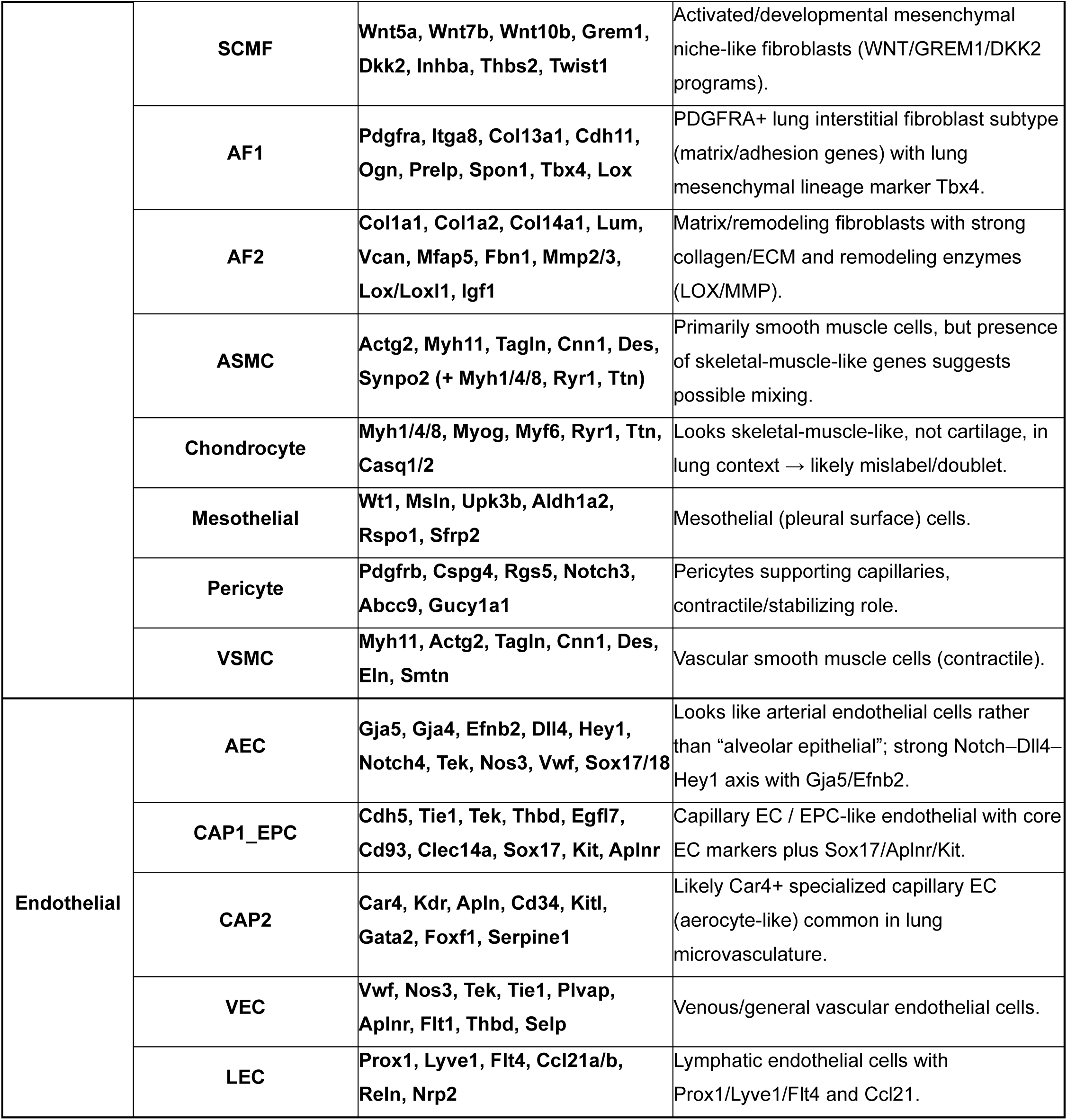
Annotated cell types and representative marker genes. Mouse Lung Cell Reference v1.1 labels, major cellular lineages, canonical marker genes, and final cell-type annotations used for downstream Xenium spatial analyses are shown.

**Table S2.**
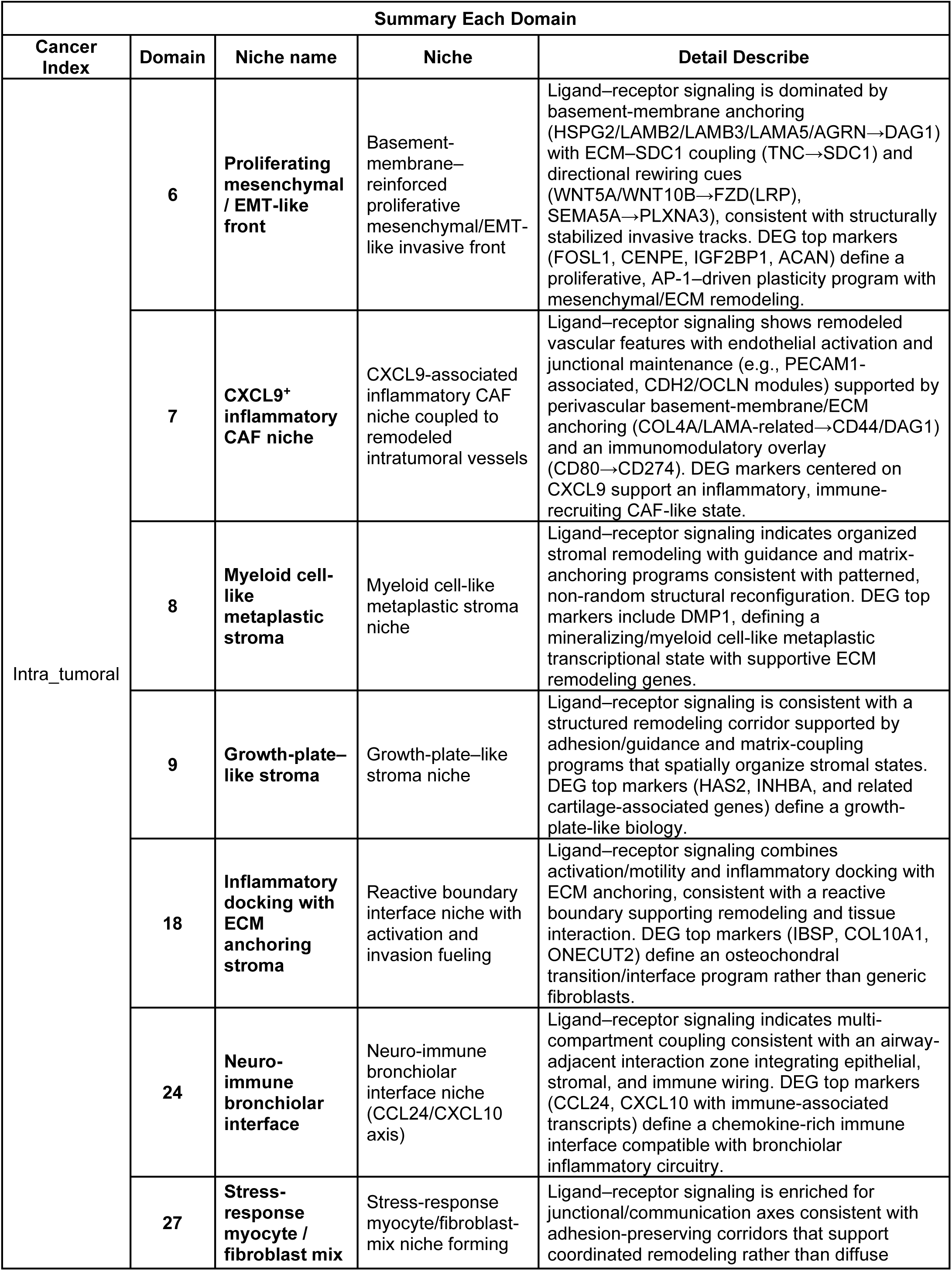
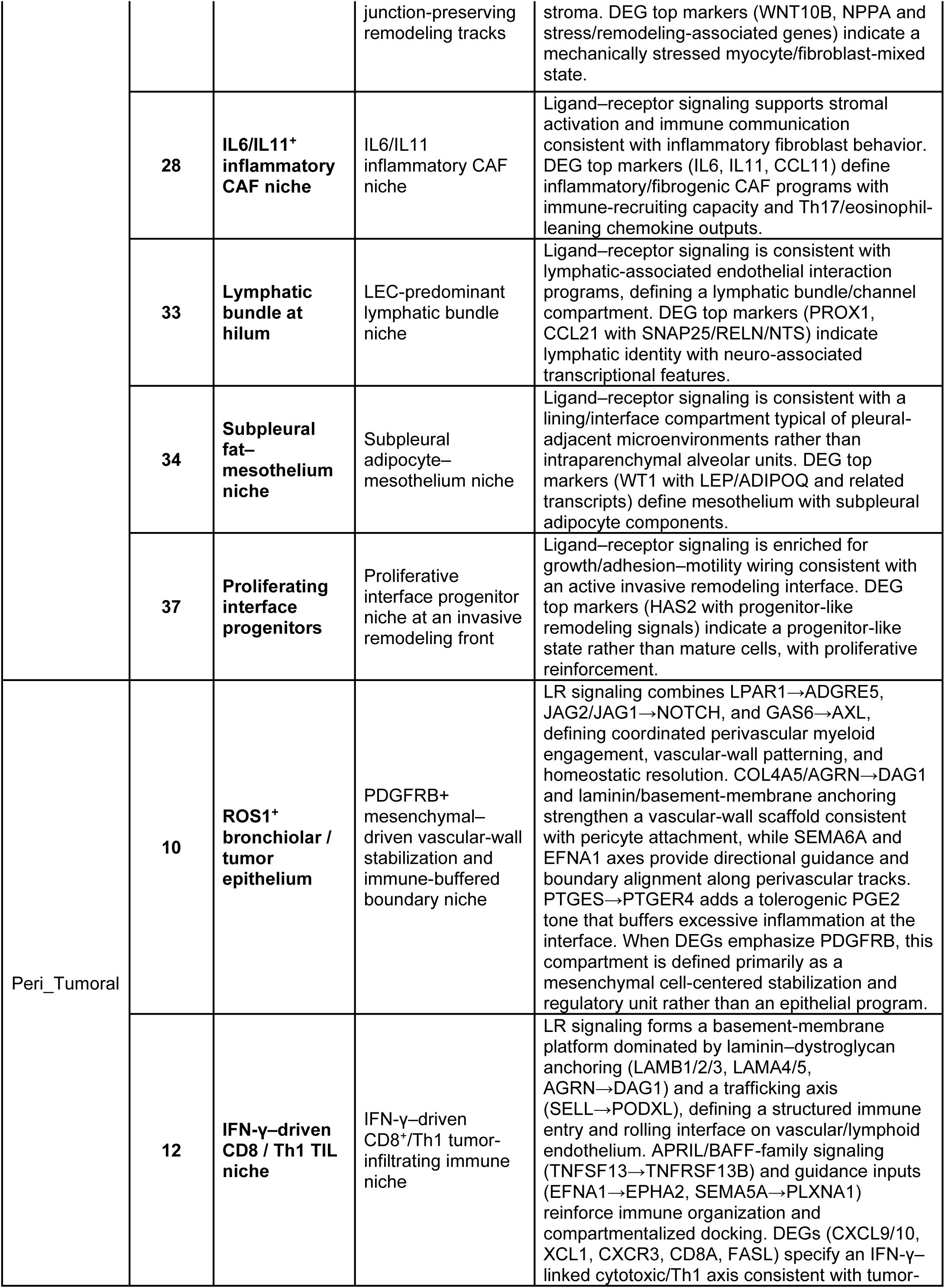
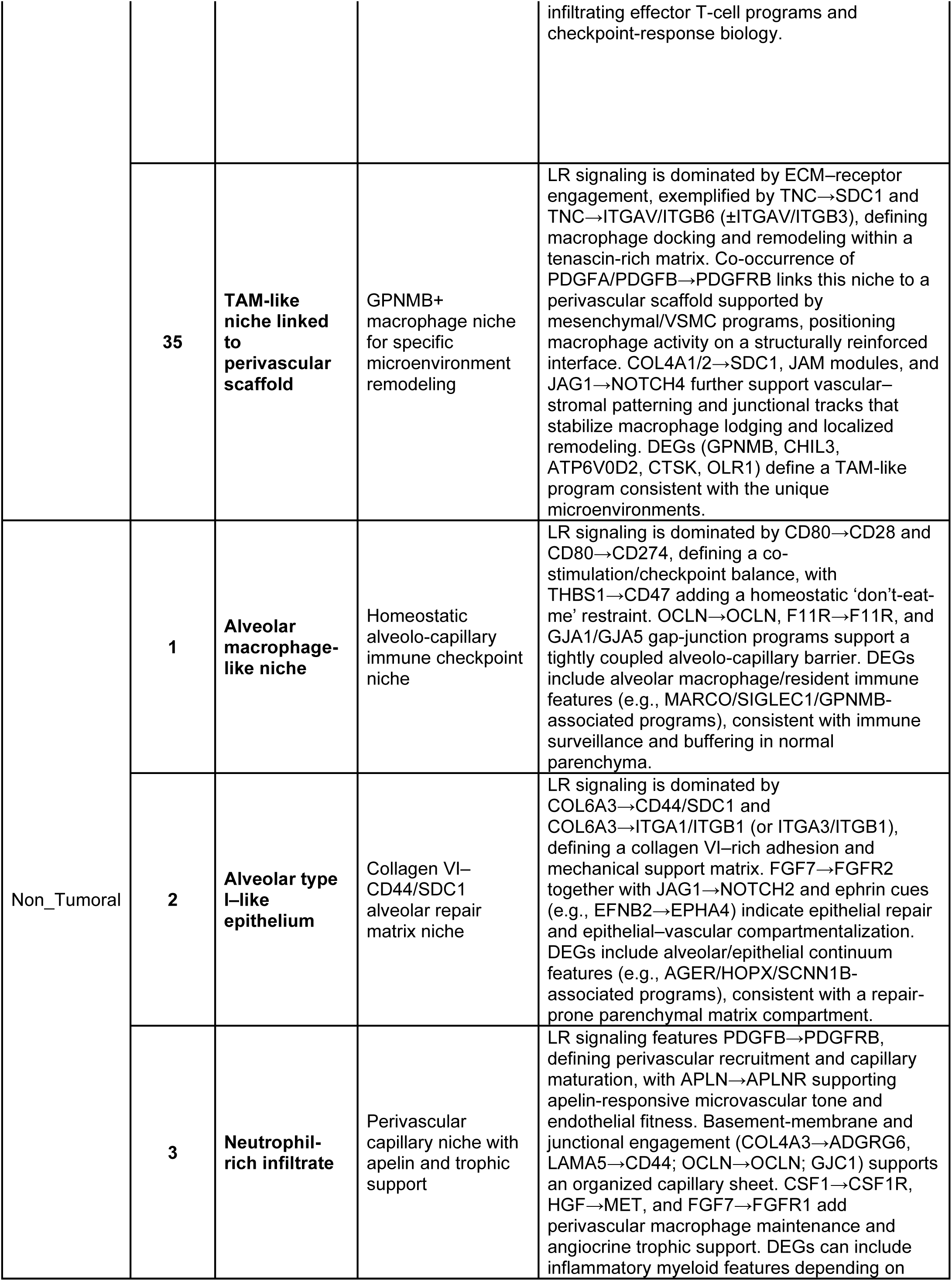
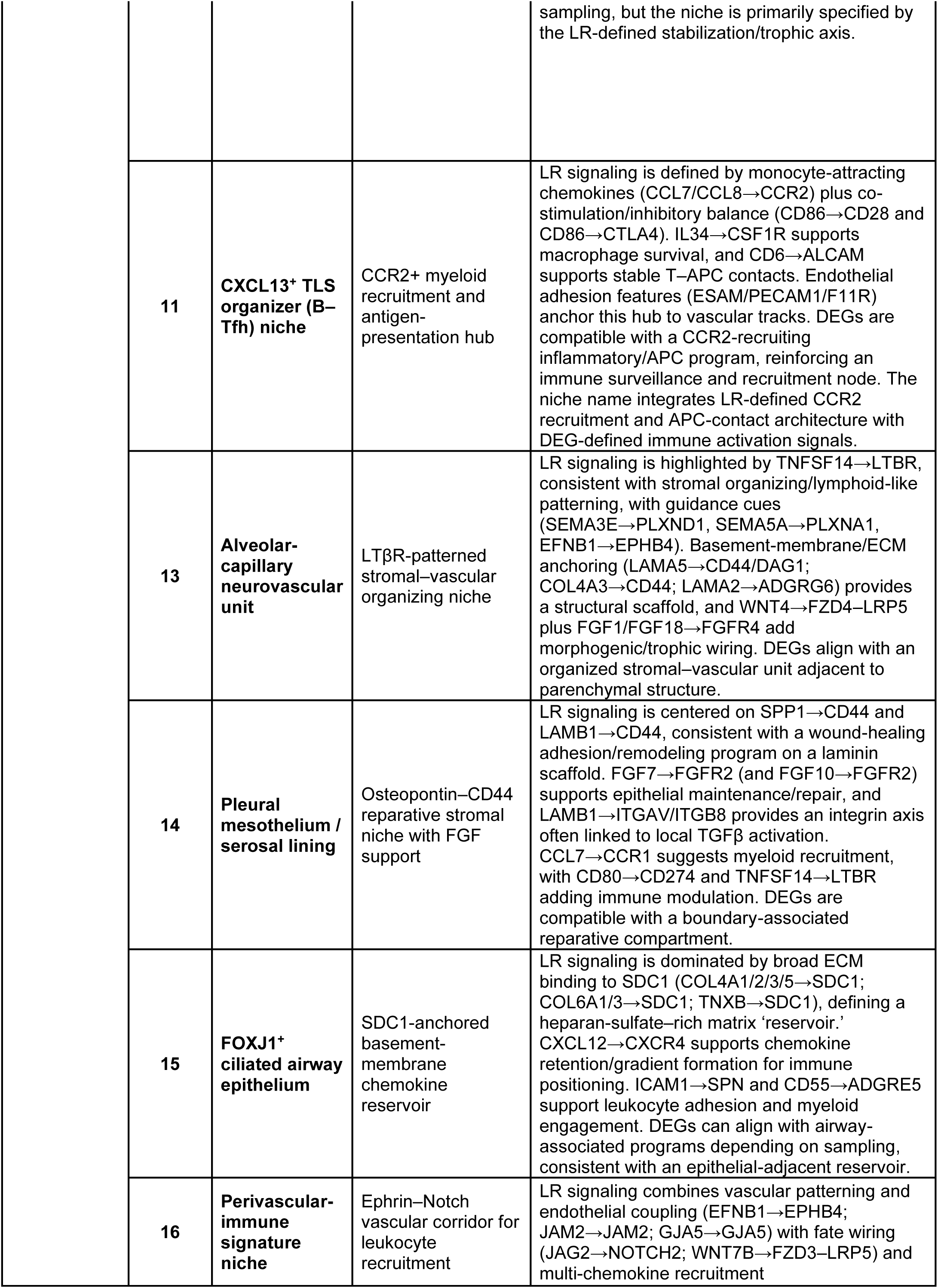
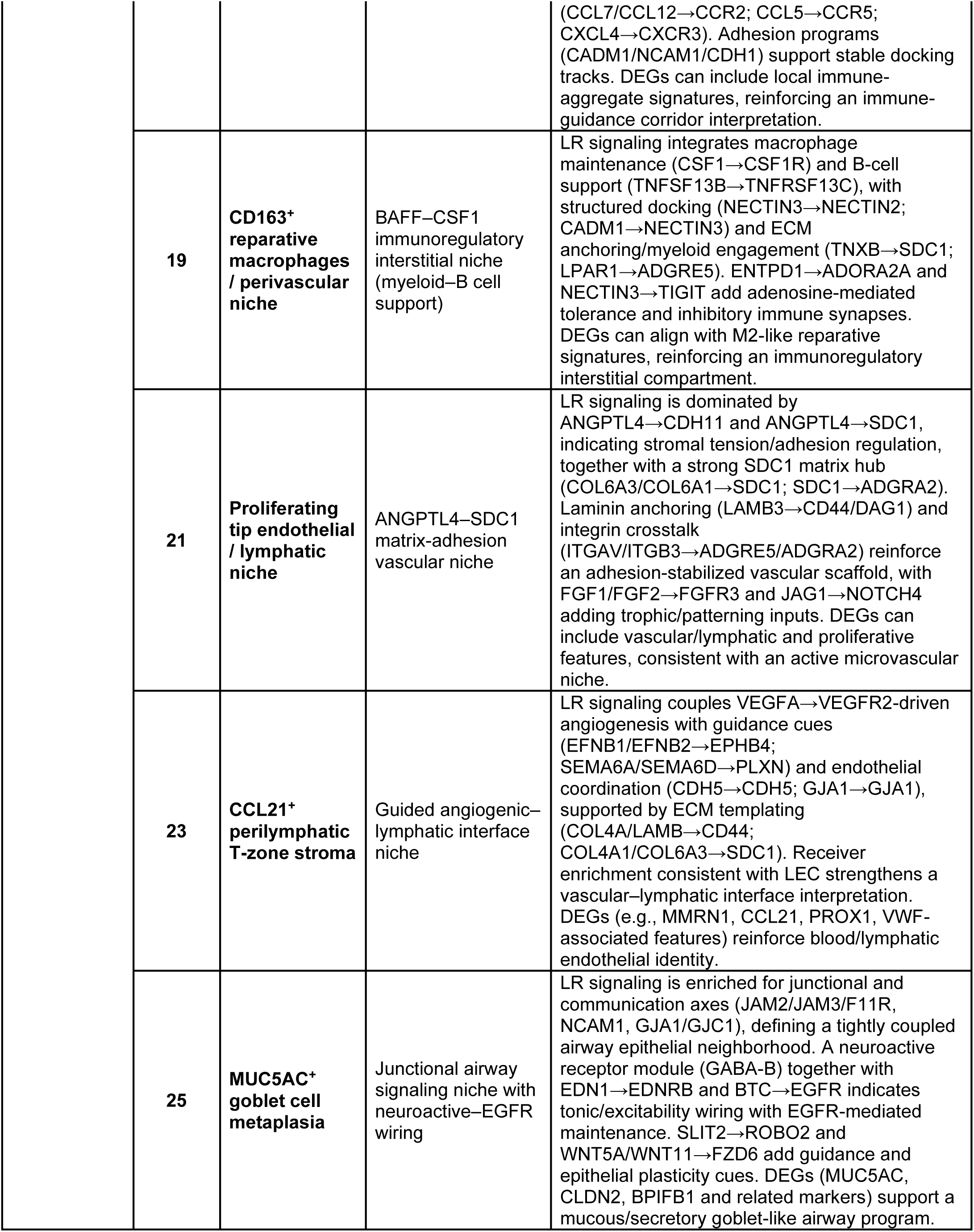
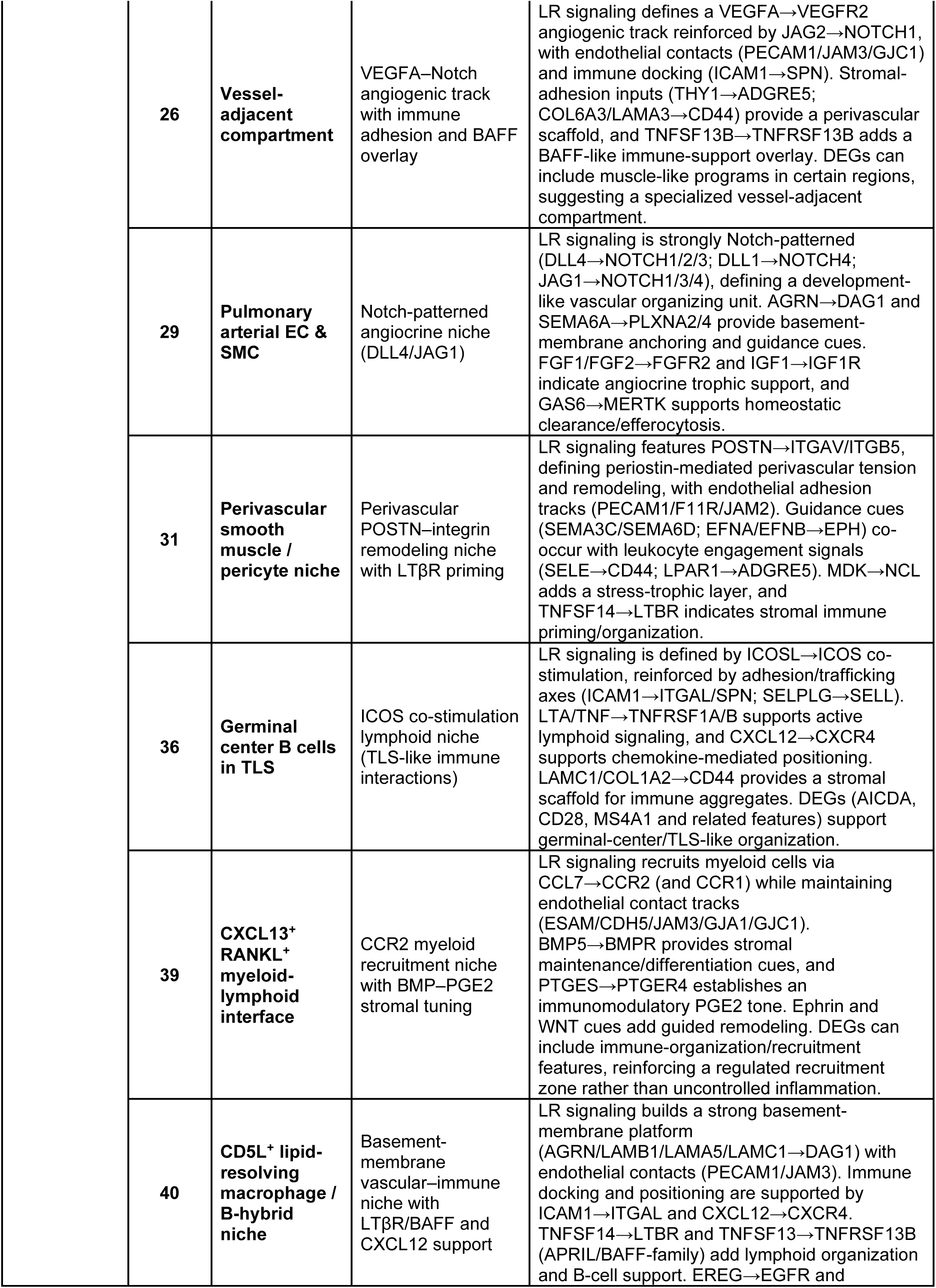
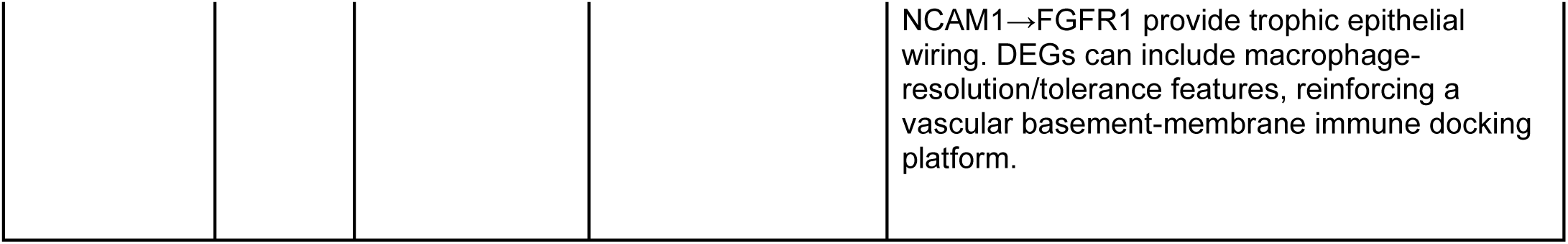
Cellular, molecular, and spatial characteristics of BANKSY-defined domains in the murine lung cancer ecosystem. Comprehensive summary of the spatial domains identified by BANKSY clustering in the murine lung cancer Xenium dataset. For each domain, the table presents its cellular composition, representative marker genes, spatial distribution, transcriptional characteristics, Cancer Index–based classification, and final biological annotation. Spatial domains were organized into intra-tumoral, peri-tumoral, and non-tumoral compartments and were further assigned to representative anatomical structures, tumor-associated micro-niches, or tumor nodule types where applicable. These annotations provide the basis for the structural, functional, and intercellular communication analyses described throughout the manuscript.

**Table S3.**
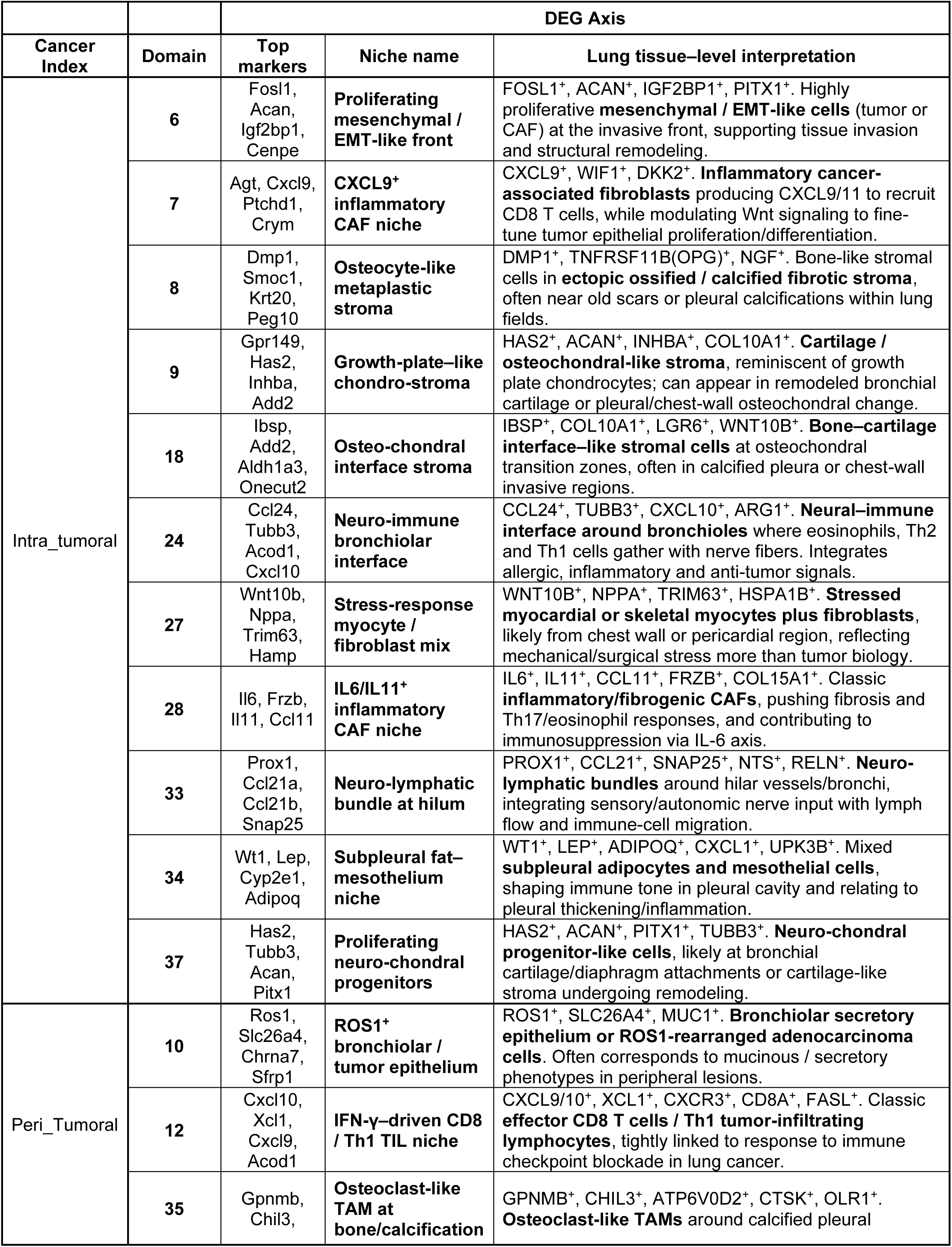
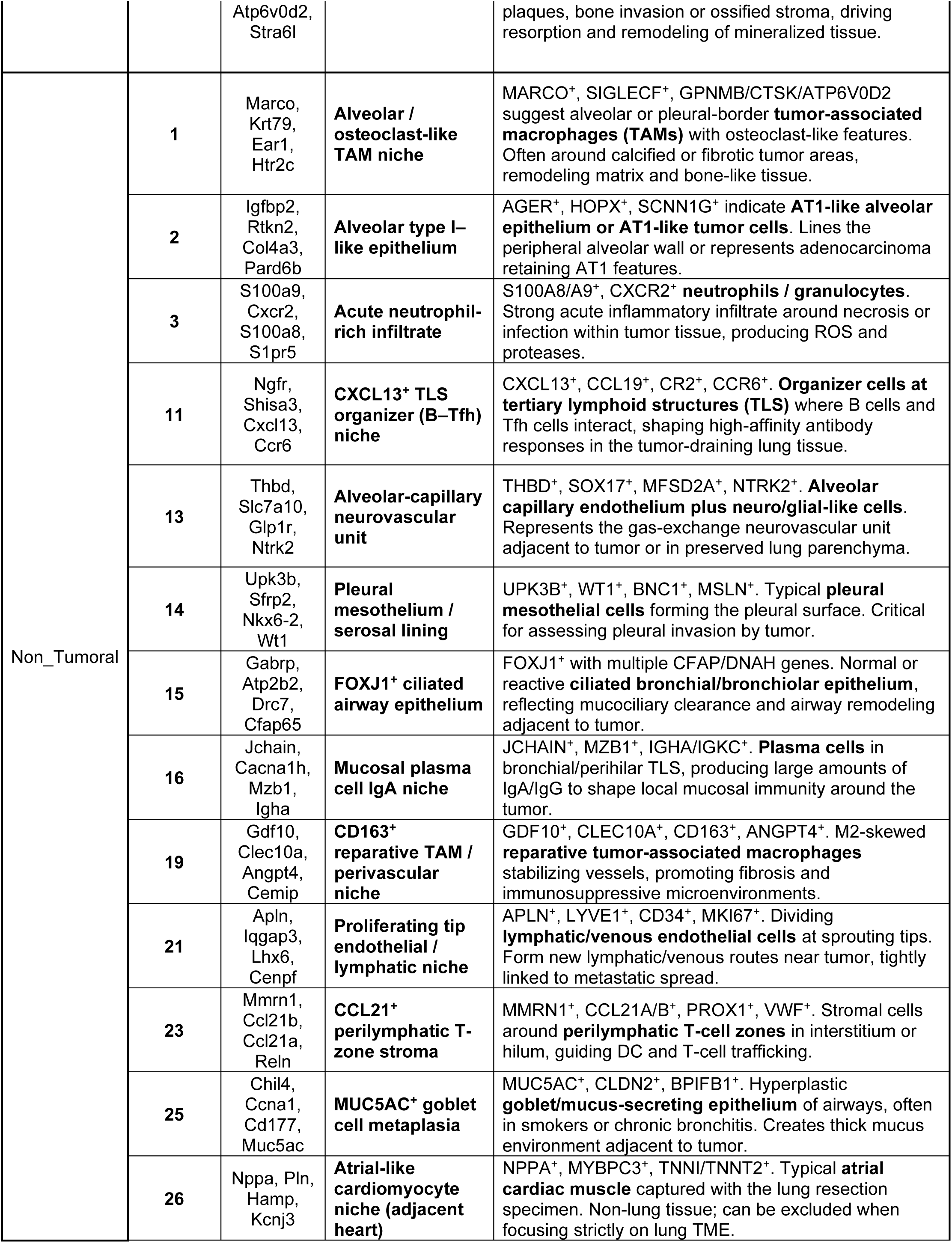
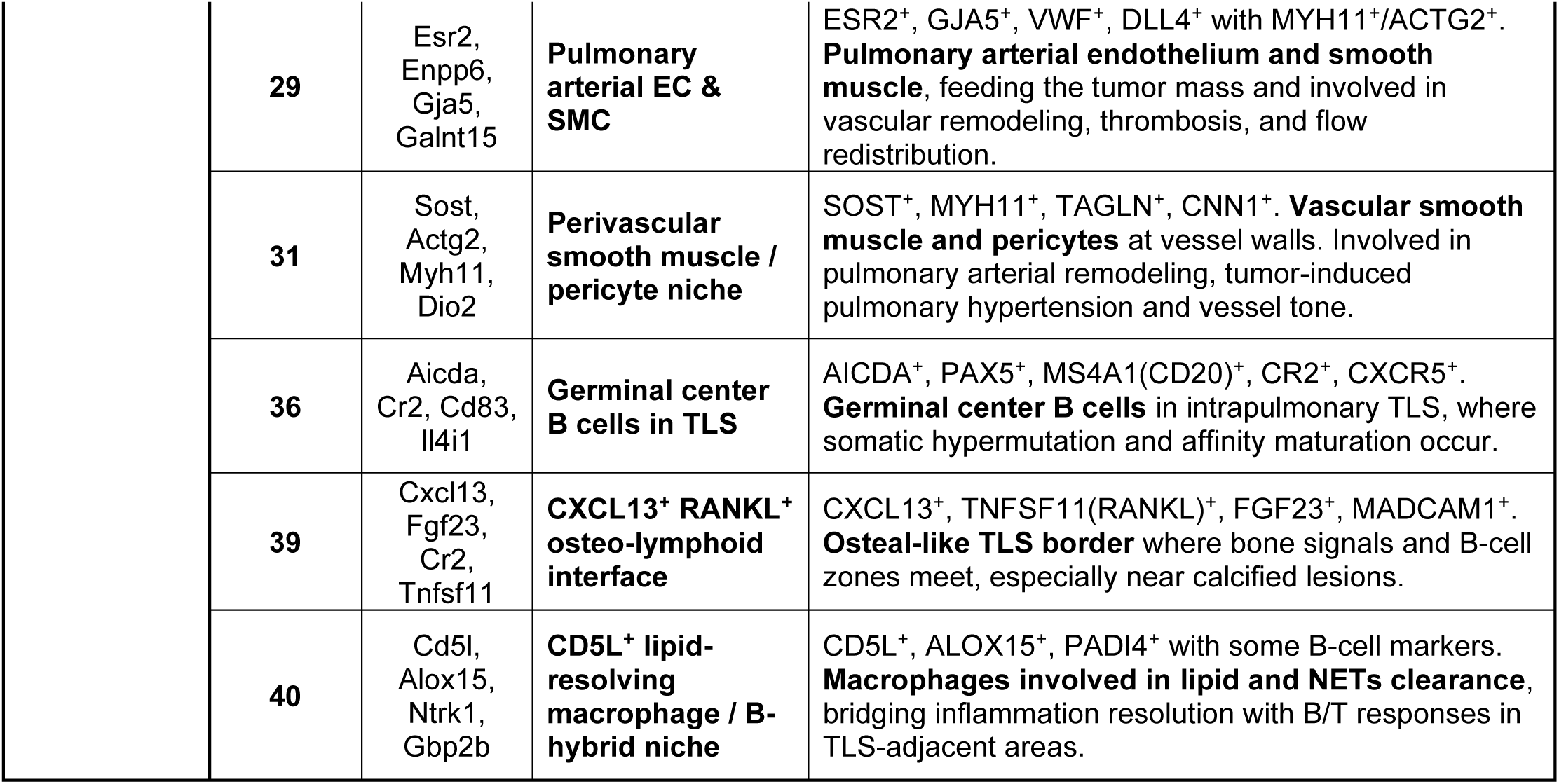
Differential gene expression and molecular characteristics of spatial domains and multicellular micro-niches. Summary of differentially expressed genes and molecular features associated with the spatial domains and multicellular micro-niches examined in this study. The table includes domain-enriched genes, representative lineage and functional markers, transcriptional programs, and statistical results from comparisons between biologically related spatial domains. These data support the molecular characterization of tumor-associated, peri-tumoral, and non-tumoral niches and provide additional information for the interpretation of the spatial domain identities and functional states presented in the main figures.

